# Sub-retinal transplantation of human iPSC-derived retinal sheets: A promising approach for the treatment of macular degeneration

**DOI:** 10.1101/2023.10.19.562811

**Authors:** Andrea Barabino, Katia Mellal, Rimi Hamam, Anna Polosa, May Griffith, Jean-François Bouchard, Ananda Kalevar, Roy Hanna, Gilbert Bernier

## Abstract

Retinal degenerative diseases affect millions of people worldwide, and legal blindness is generally associated with the loss of cone photoreceptors located in the retina’s central region called the macula. Currently, there is no treatment to replace the macula. Addressing this unmet need, we employed control isogenic and hypoimmunogenic induced pluripotent stem cell (iPSC) lines to generate spontaneously polarized retinal sheets (RSs). They presented the advantage of facile customization and large-scale production, exhibiting a polarized 3D architecture within a 2D cell culture environment, thus readily adaptable for clinical applications. RSs were enriched in retinal progenitor and cone precursor cells, which could differentiate into mature S- and M/L-cones in long-term cultures. Single-cell RNA-seq analysis showed that RSs recapitulate the ontogeny of the developing human retina and are devoid of pluripotent stem cells. Isolation of neural rosettes for sub-retinal transplantation effectively eliminated unwanted cells such as RPE cells. In a porcine model of chemically induced retinal degeneration, grafts integrated the host retina and formed a new, yet immature, photoreceptor layer, exhibiting viability for up to 2 months. In one transplanted animal that met all criteria, including correct apicobasal orientation and seamless graft integration, functional and immunohistochemical assays suggest that grafts exhibited responsiveness to light stimuli and established putative synaptic connections with host bipolar neurons. This study underscores the potential and challenges of iPSC-derived RS for clinical applications while establishing the optimal methodology and conditions for RS transplantation, thus paving the path for future long-term functional investigations using RS grafts.

## MAIN

In most retinal degenerative diseases, loss of visual function results from the death of photoreceptors, the specialized light-sensitive cells involved in phototransduction^1^. Photoreceptors exist in two types, rods and cones. Rods are the most abundant photoreceptors in the human retina and respond to dim light. They are involved in night vision and important for peripheral vision. Cones respond to intense light and are required for color, daylight, and high-resolution central vision ^2,3^. In contrast to other mammals, the eye of modern primates (apes, monkeys, and hominins) contains a unique circular structure of 4-5 mm of diameter called the macula located near the center of the retina ^4,5^. The macula is highly enriched in cone photoreceptors and has a cone-only smaller region of ∼1.5 mm in diameter called the fovea ^4,5^. Retinal degenerative conditions (late-stage retinitis pigmentosa, macular degenerations, and inherited retinal dystrophies) all share the common features of loss of cone photoreceptors and central (macular) vision ^2^. Retinitis pigmentosa is one of the most common and genetically heterogeneous retinal degenerative diseases ^6^. While the disease primarily affects rods, it usually progresses toward legal blindness when cones located in the center of the retina are lost. Loss of central vision is also predominant in cone-specific diseases such as Stargardt disease, cone dystrophies, cone/rod dystrophy, Leber congenital amaurosis (LCA), and macular degeneration^7^.

Despite its major impact on quality of life, therapeutic interventions against vision loss remain of limited effectiveness and are confined to a small number of cases. Gene therapy approaches aiming at correcting or silencing the disease-causing mutation or replacing the defective gene are promising avenues. They rely on the injection of viruses and have, to date, demonstrated limited success, primarily restricted to specific cases, such as patients with LCA resulting from a mutation in RPE65^8^. Gene therapy is also limited because it focuses on a single gene or mutation at a time when RDs are associated with mutations in hundreds of genes^9,10^. Hence, while gene therapy can prevent photoreceptor degeneration, it cannot replace lost photoreceptor cells. Prosthetic sub-retinal implants (e.g., Argus II) have also been developed. However, these devices are invasive and can only provide limited visual stimuli in patients ^11^. Because human vision depends mainly on the macula for most activities, there is a need for cone cell replacement therapy to restore central vision.

Although promising, photoreceptor transplantation therapy in animal models has been debated for its efficiency and suitability. It has been suggested that grafted rod precursors can undergo material transfer (RNA, protein, and exosomes) with the host photoreceptors, potentially leading to non-cell-autonomous effects ^12,13^. On the other hand, work from other groups suggests that dissociated human photoreceptors grafted in the sub-retinal space can integrate and differentiate efficiently into the photoreceptor cell layer of immunodeficient mice, and extensive analyses confirmed that material transfer or cell fusion does not occur in this context ^14^. Notably, when grafted into the retina of late-stage *rd1* mice lacking photoreceptors, mouse iPSC-derived retinal tissues or dissociated human iPSC-derived cone precursors can form presumptive synapses with endogenous bipolar neurons and improve visual function, thus also generally excluding the thesis of material transfer to endogenous photoreceptors^15–17^. Results from these important experiments also suggest that even in congenitally blind mice, the central connections from the retina are still relatively intact and functional ^15,17,18^. At last, while pioneer work from Sasai’s laboratory provided a revolutionizing method to generate human retinal organoids ^19–22^, their heterogeneous nature may represent an obstacle to retinal transplantation therapy, such as the formation of neural rosette-like structure within the host sub-retinal space^17,23^. Retinal sheets from fetal retina have shown connectivity and visual recovery in a rat model of retina degeneration^24,25^ and retinal organoids have recently been employed to produce sheet-like grafts for transplantation in rats, yielding promising outcomes, but also developing neural rosette-like structures within the host sub-retinal space ^26^. Thus, the current protocols to purify dissociated cone precursors or to generate sheet-like grafts from retinal organoids involve laborious and specialized manual procedures, resulting in limited quantities and scalability for human clinical application.

We previously reported on an independent method to produce abundant cone precursors from human embryonic stem cells (ESCs) and iPSCs, forming uniform and polarized retinal sheets (RSs) with a 3D architecture in a 2D culture environment. This feature allows for easy customization by cutting the material into the desired size and shape for grafting, providing ample material for multiple transplants ^27,28^. The method relies on the use of human recombinant COCO (also called DAND5), a Cerberus-like family member protein that can antagonize soluble BMP, TGFβ, and WNT ligands. The method allows the spontaneous formation of a ∼100µm thick, multilayered, and polarized RS containing up to ∼60% of cone progenitors and precursors, thus possibly suitable for macula replacement therapy^27^.

We herein describe the molecular and cellular characterization of RSs produced from isogenic control and hypoimmunogenic universal donor (UDC) iPSC lines. Using a cobalt chloride-induced model of severe photoreceptor degeneration in the Yucatan minipig, we show that RS punches transplanted in the sub-retinal space can form a new but yet immature cone photoreceptor cell layer within the host retina, with evidence of visual response to bright light at the lesion site.

## RESULTS

### Single-cell RNA sequencing reveals cell population dynamics during retinal sheet development

We induced the differentiation of the isogenic WTC/Actin-GFP (WTC^Act^), WTC/Tubulin-RFP (WTC^Tub^), and WTC-UDC (WTC^UDC^) iPSC lines into RSs (**Fig. S1A, and Material and Methods**). CRX is a master regulator of photoreceptor development, and flow cytometry analysis at DIV60 revealed that when compared to unstained control cells, ∼65% of the cells were positive for CRX (12% CRX-high and 53% CRX-low) (**Fig. S1B**). By RNA-sequencing (RNA-seq), global gene expression of DIV60 RSs was compared to that of the human embryonic retinal development atlas ^29^. This revealed that RSs express the pan photoreceptor genes *CRX*, *OTX2*, and *ROM1*, and the cone-specific genes *OPN1SW* (encoding S-opsin) and *PDE6B* (encoding for a light-responsive phosphodiesterase) at levels that matched day 67-94 of the human embryonic retina. (**Fig. S1C**). To test the functionality of RSs *in vitro*, we measured the proportion of endogenous cGMP degraded by light-responsive phosphodiesterases. This revealed light-induced cGMP degradation in RSs when compared with those treated with a phosphodiesterase inhibitor, thus showing a biochemical response to bright light (**Fig. S1D**) ^30–34^.

To deepen the characterization of the cell populations, we performed single-cell RNA-seq (scRNA-seq). DIV60 RSs were found to contain about all cell types present in the human embryonic eye (control human retina samples ranging from day 94 to day 129 post-conception)^35^, except for gene sets specific to bipolar neurons, rod photoreceptors, astrocytes, and Müller glia (**Fig. 1A**). UMAP analysis revealed that retinal progenitor cells (RPCs) were the most abundant cell types present in RSs (**Fig. 1A**). Developmental kinetics analysis of RSs at DIV5, DIV30, and DIV60 revealed a linear relationship in pseudo-time representation (**Fig. 1B**), with the identification of markers for early (*MKI67* and *PAX2*), intermediate (*ASCL1* and *VSX2*) and late (*RCVRN* and *RPE65*) groups (**Fig. 1C**). *RAX, PAX6, LHX2, SIX3, SIX6,* and *VSX2* are expressed at the earliest stages of retinal development ^36–44^ and delineated a prominent RPC population present in RSs at DIV5, DIV30, and DIV60 (**Figs. 1D and S2A**).

**Figure 1.**
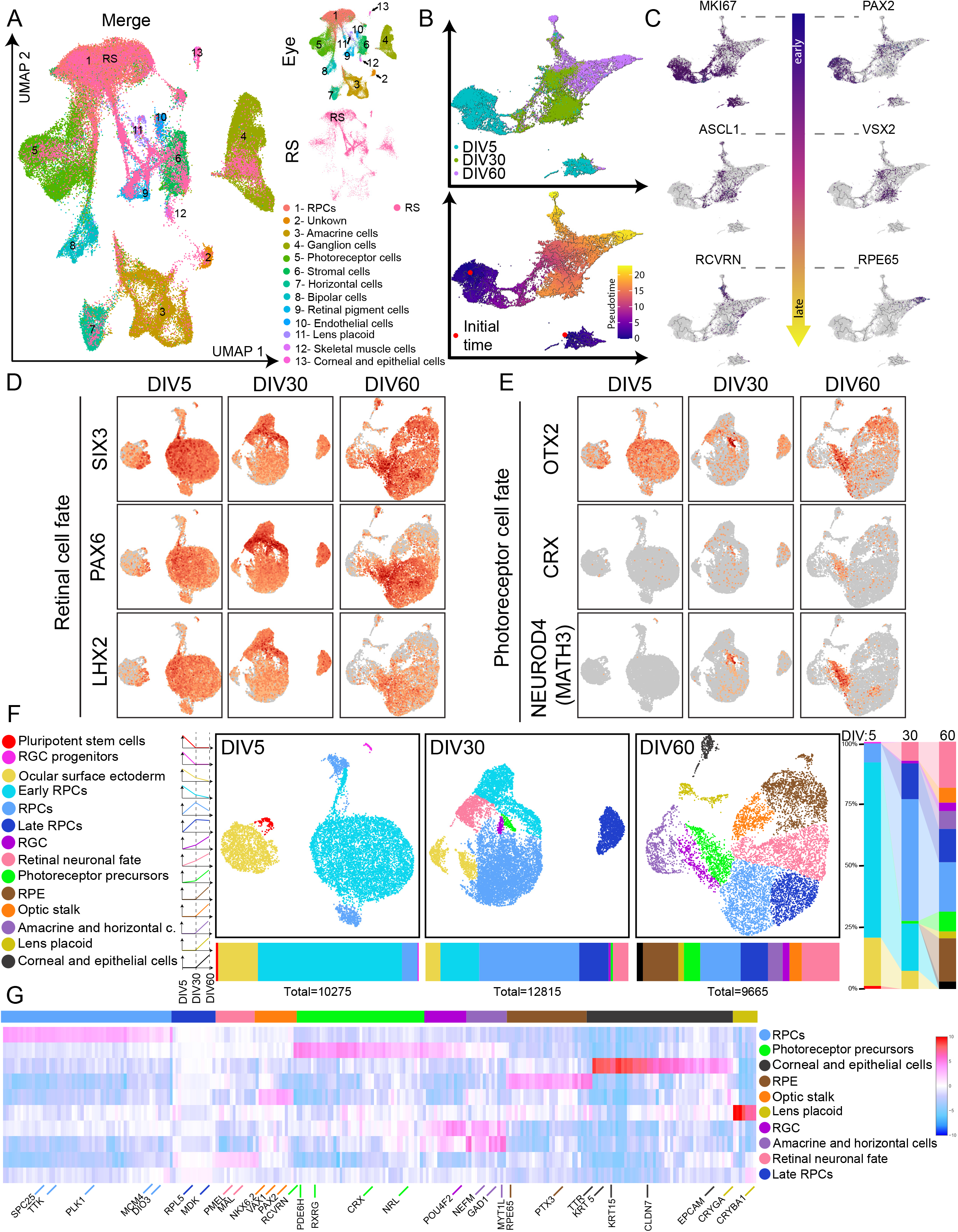
Identification of cell populations and developmental kinetics of retinal sheets using single-cell RNA sequencing. A-UMAP distribution of the single cell RNA-seq from the embryonic human eye and the retinal sheet DIV60 distributed on the same 2D space. The embryonic human eye dataset is from the Descartes database. The labels inferred for each group of cells are deduced from the classifier of the Descartes dataset. B-Pseudotime analysis on scRNA sequencing using Monocle3 on three different time points of our differentiation DIV5, DIV30, and DIV60, which refer to 5, 30, and 60 days after the beginning of differentiation. Each point represents the transcriptome in a single cell. The calculated cells scattered on the UMAP graph generally preserved the grouping of the differentiation stages. The pseudotime calculated by the monocle3 also reflected the biological time of the differentiation stages, with the earliest being the DIV5 and the most advanced being the DIV60. C-Featureplot representation on the pseudotime UMAP graph highlighting early expressed markers: MKI67 and PAX2. Retinal fate markers: ASCL1 and VSX2. And finally, terminally differentiated markers: RCVRN and RPE65. Each point represents the transcription levels of the gene in a single cell. D-Featureplot representation plotted on the UMAP graphs of the DIV5, DIV30, or DIV60 of genes that highlight the retinal cell fate: SIX3, PAX6, and LHX2. Each point represents the transcription levels of the gene in a single cell. E-Featureplot representation plotted on the UMAP graphs of the DIV5, DIV30, or DIV60 genes that highlight the photoreceptor cell fate: OTX2, CRX, and NEUROD4. Each point represents the transcription levels of the gene in a single cell. F-UMAP graphs identify the cell types in each differentiation stage; the graph is color-coded according to the cell types. Each point represents a single cell. Under each graph, a horizontal slice graph is presented to show the percentage of each cell type in the corresponding differentiation stage, along with the total number of cells analyzed for the before-mentioned differentiation stage. On the right, a vertical slice shows the evolution of the culture composition in terms of cell lineage. G-Heatmap of the most differentially expressed genes between the cell types at DIV60. Each row represents a cell type, and each column represents a distinct gene. These genes can be grouped according to the cell type in which they are mostly expressed, as shown with the color-coded bar above the heatmap. The colors follow the same code as in F. We highlighted some genes that would help to identify and confirm the group identity.

OTX2 and CRX are closely related transcription factors critical for photoreceptor development. OTX2 is expressed early in retinal development, including in retinal progenitor cells, photoreceptor precursors, and mature photoreceptors. In contrast, CRX expression begins later, becoming prominent only as retinal progenitor cells differentiate into photoreceptor precursors and mature photoreceptors ^45–50^. Both OTX2 and CRX are also expressed in a subset of bipolar neurons. Accordingly, *OTX2* was expressed earlier and more broadly than *CRX* and all other photoreceptor markers in RSs at DIV5, DIV30, and DIV60 (**Figs. 1E and S2B**). Although *OTX2* is expressed in all photoreceptor lineage cells, it is also found in the retinal pigment epithelium (RPE), in contrast with CRX. Our annotation of cell types in RSs at DIV5, DIV30, and DIV60 revealed that a significant fraction of *OTX2*-expressing cells at DIV60 were indeed RPE cells (**Figs. 1E-F**). This also revealed the presence of two main cell populations at DIV5, i.e., RPCs and ocular surface ectoderm. The ocular surface ectoderm can give rise to both the lens and cornea. This cell population expressed Keratin genes as well as *PAX6* and *SIX3*, both required for ocular surface ectoderm specification in mice (**Figs. 1F and S2D-E**) ^51–56^.

Rare putative pluripotent stem cells expressing both *POU5F1* and *NANOG* at DIV5 were found very close to the ocular surface ectoderm cell population but distant from RPCs (**Figs. 1F, S2D-E and S3A-B**). Between DIV5 and DIV30, the early RPC population (expressing high levels of *MKI67* and *PCNA*) gradually diminished and disappeared by DIV60, making room for RPCs, late RPCs, photoreceptor precursors, and a distinct retinal precursor cell population classified here as retinal neuronal fate (**Figs. 1F and S2D-E**). Notably, putative pluripotent stem cells were not observed at DIV30 and DIV60, consistently with the absence of teratoma formation in NOD/SCID mice grafted with dissociated DIV60 RSs (**Figs. 1F, S2D-E and S3A-C**). Heat map representation of annotated genes in DIV60 RSs further revealed key markers distinguishing the identified cell populations, including lens placoid (*CRYGA* and *CRYBA1*) and corneal cells (*KRT5, KRT15,* and *CLDN7*) (**Fig. 1G**)^57–60^. Pax2 expression in early mouse eye development occurs in the optic stalk and marks the boundary between the presumptive neural retina (Pax6+) and the optic nerve head (Pax2+) ^61^. The *PAX2*, *VAX1,* and *NKX6-2-*positive group thus represents a defined ocular neuroectoderm cell population corresponding to the presumptive optic stalk and later to its derivatives, such as the optic nerve and optic nerve head (**Figs. 1G and S2B**)^62,63^.

In each cluster, we observe notable cellular diversity. For example, late RPCs, while retaining their characteristic identity, also contain a distinct subgroup (6.4%) that begins expressing *RCVRN*, signifying a transition towards precursor-like states (**Figs. 2B-C and S2C**). Similarly, in the precursor cell population, many cells still retain expression of *PAX6* and *VSX2* (27.8%), signifying their persistence in a more immature state (**Figs. 1D and S2A**). Intriguingly, a subset of precursor cells displays a shift in identity by expressing *PDE6H, GNGT1, GNGT2,* and *ARR3* suggesting a progression toward a more mature cellular state (**Fig. S2C**)^64^. These observations emphasize the dynamic nature of cell identities within clusters and highlight the possibility of transitional states within specific cell populations.

**Figure 2.**
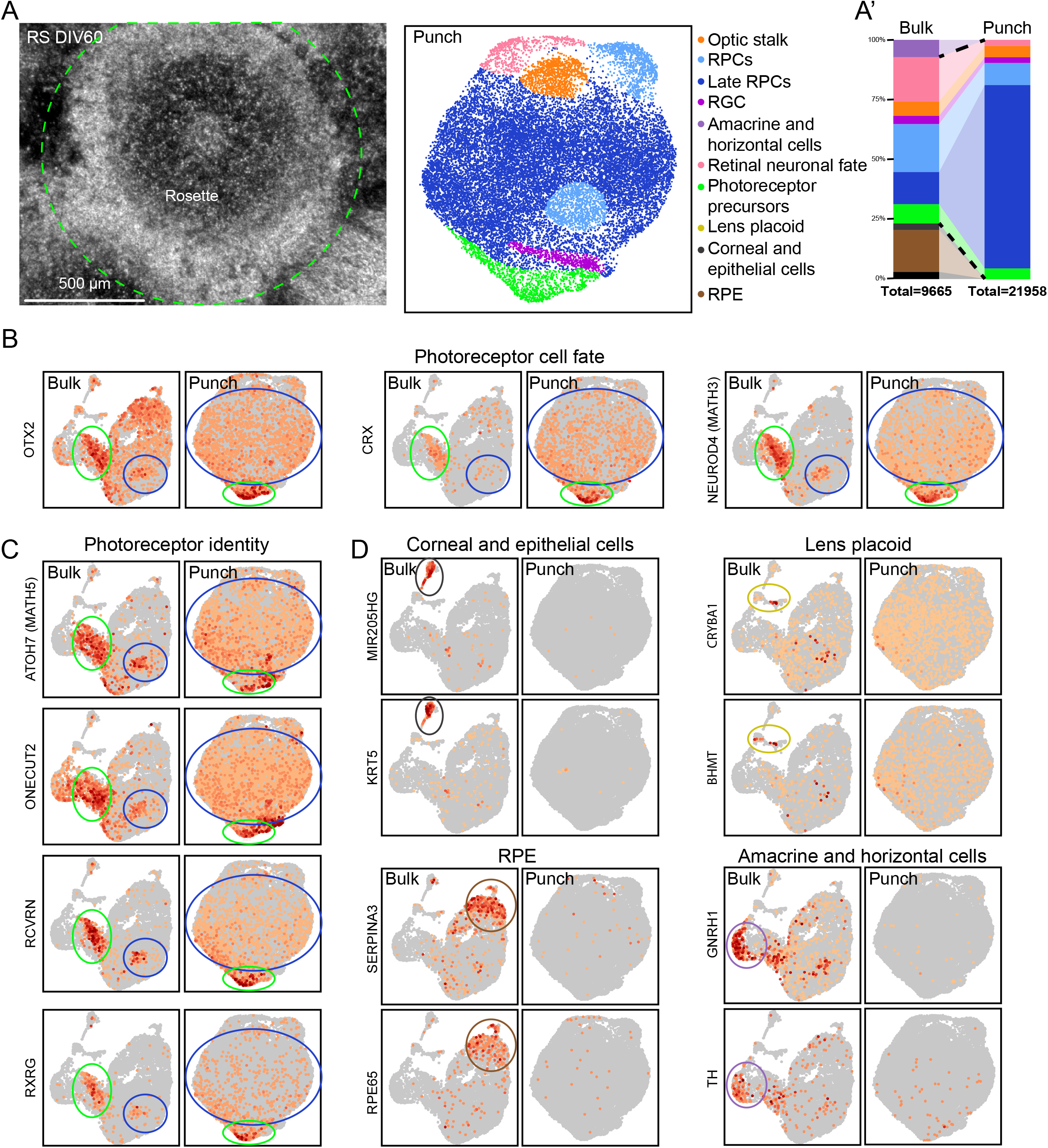
Characterization of large neural rosettes by single-cell RNA-sequencing. A-DIC image of retinal rosettes at 60 days of differentiation. One of the rosettes is highlighted with a green circle to illustrate. On the right, the UMAP graphs identify the cell types in the punch from RS culture; the graph is color-coded according to the cell types. Each point represents a single cell. On the left (A’), a vertical slice graph shows the percentage of each cell type in the bulk population versus the punch population and the total number of cells analyzed for the above-mentioned populations. The dashed line shows the cell populations of the bulk RS cultures at DIV60 that were preserved in the punch. B-Featureplot representation plotted on the UMAP graphs of the DIV60 or the punch of genes that highlight the photoreceptor cell fate. Each point represents the transcription levels of the gene in a single cell. These graphs show the preservation of the photoreceptor cell population in the punch. A circle color-coded with the cell type color highlights the position of these cell populations. C-Featureplot representation plotted on the UMAP graphs of the DIV60 or the punch of genes that highlight the photoreceptor identity. Each point represents the transcription levels of the gene in a single cell. These graphs show the preservation of the photoreceptor cell population in the punch. A circle color-coded with the cell type color highlights the position of these cell populations. D-Featureplot representation plotted on the UMAP graphs of the bulk DIV60 RS or the punch of genes that highlight the lost cell populations: Keratocytes, Lens placoid, RPE, amacrine, and horizontal cells. Each point represents the transcription levels of the gene in a single cell. These graphs show the loss of these cell populations in the punch. A circle color-coded with the cell type color highlights the position of these cell populations.

### Large neural rosettes are enriched for photoreceptor precursors and RPCs

While RSs are enriched for RPCs and photoreceptor precursors, they also contain other cell types that could interfere with their application in therapy. Large neural rosettes were isolated using a biopsy punch and analyzed by scRNA-seq (**Fig. 2A**). When compared to DIV60 RS cultures (i.e., the bulk population), RS punches were highly enriched for the late RPCs cluster, with about the same proportion of photoreceptor precursors (**Figs. 2A’-C and S4A-E**). Notably, RS punches exhibited a significant depletion of RPE, amacrine, horizontal, lens placoid, and corneal epithelium cells (**Fig. 2D**). Consequently, this resulted in a highly enriched population of late RPCs and photoreceptor precursor cells suitable for transplantation therapy.

We next conducted immunofluorescence (IF) analyses on retinal sheets at various time points throughout the differentiation protocol (**Fig. 3A**). DIV60 RS punches were composed of a layer of 4-8 nuclei expressing CRX, OTX2, VSX2, and RCVRN (**Figs. 3B and S4F**). The tissue was polarized, with RCVRN and PNA labeling being present at the apical side over the nuclear layer (**Figs. 3B and S5A**). The photoreceptor precursor nuclei on the apical side were strongly positive for CRX and OTX2, with the nuclei localized in the tissue’s center and the basal side positive for VSX2 (**Figs. 3B and S5A**). In RS punches from both WTC^UDC^ and WTC^Act^ cell lines, 20-25% of the cells show strong expression of CRX and OTX2 (**Fig. 3C**). These findings align with the scRNA-seq results and are comparable to FACS analysis of the bulk population when focusing on CRX-high cells (**Figs. 2B and S1B**).

**Figure 3.**
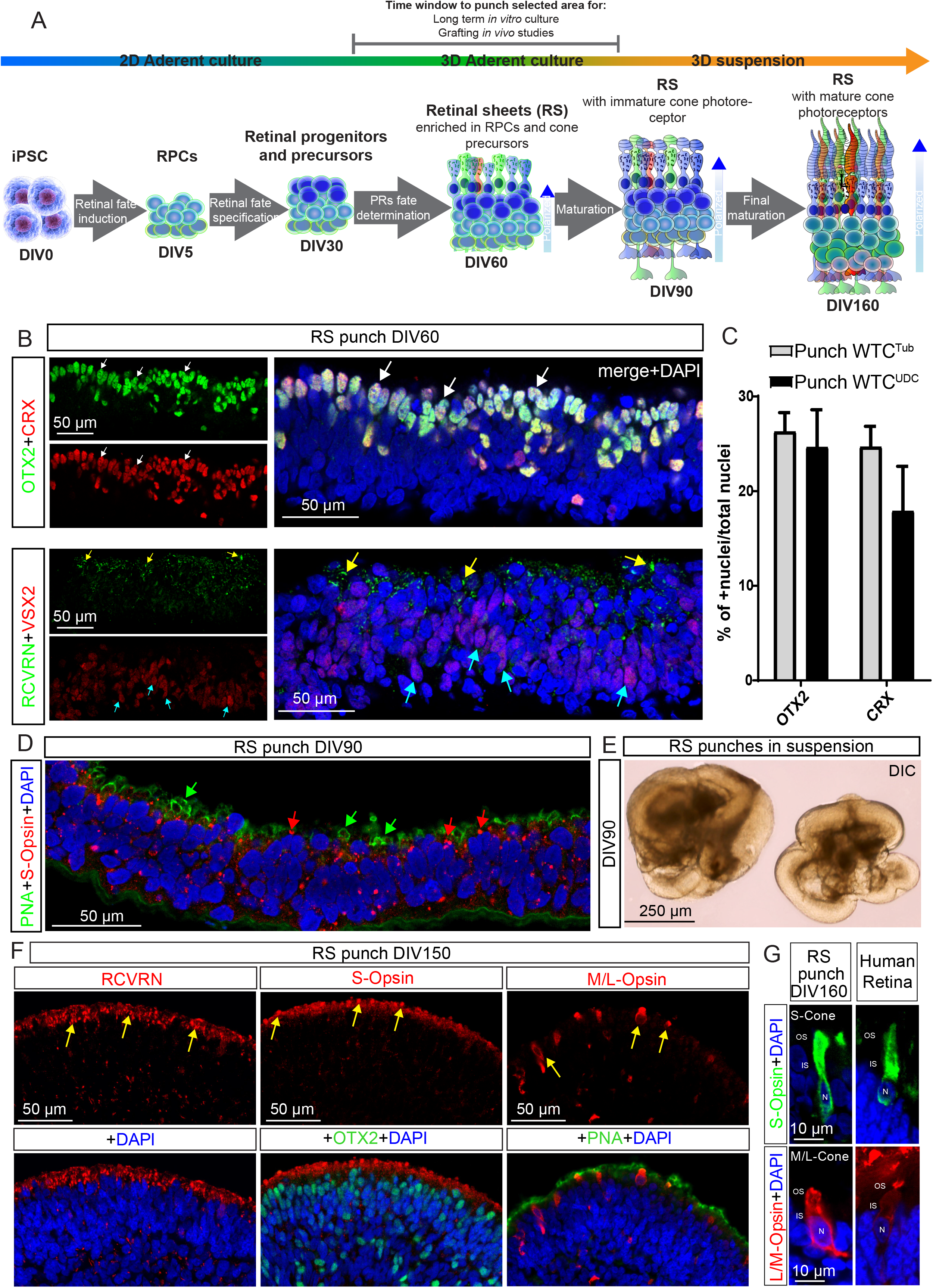
Histological characterization of neural rosettes and retinal organoids. A-Overview of the timeline of the differentiation procedure with key steps and genes/proteins characteristic of each phase identified either by scRNAseq and/or IF. B-Confocal IF images of cryosections of retinal sheet punch generated from the WTC^Tub^ iPSC line. Photoreceptors fate markers: OTX2, CRX, and RCVRN. RPCs marker: VSX2. White arrows indicate photoreceptors precursors nuclei with high expression of CRX and OTX2 and negative for VSX2. Yellow arrows show apical accumulation of RCVRN in photoreceptors precursors. Cyan arrows indicate VSX2+ RPCs. C-Quantification of CRX and OTX2 positive nuclei in RSs cryosections by IF in both WTC^Tub^ and WTC^UDC^ cell lines. D-Confocal IF images of cryosections of retinal sheet punch generated from the WTC^Tub^ iPSC line differentiated for 90 days. Photoreceptors fate markers: PNA and S-Opsin. Green arrows indicate the initial formation of a rudimentary OS (PNA+) budding from the IS. Red arrows indicate granules of S-Opsin that start localizing apically in immature cone photoreceptors. E-DIC image of WTC^UDC^ retinal sheets punches in suspension culture at day 75 of differentiation. F-Confocal IF images of cryosections of retinal sheet punches of the line WTC^UDC^ at day 150 showing expression of cone photoreceptor markers RCVRN, OTX2, S-opsin, and M/L opsin. White arrows indicate OTX2+ cone photoreceptors nuclei. Yellow arrows indicate the OS with a polarized accumulation of RCVRN, S-Opsin, and M/L-Opsin. G-Confocal IF images of cryosections of retinal sheet punches at day 160 show higher magnification images of mature cone photoreceptors with elongated OS, compared with adult human cones. B, C, D, E, F, G Representative of n□=□10, 3, 3, 4, 4, 4 independent experiments, respectively. Scale bars are indicated in each figure.

Transmission electronic microscopy analysis at DIV60 further revealed the presence of tight junctions at the border of the mitochondria-rich nascent inner segments (**Fig. S5B**). Dense organelles corresponding to the basal body of primary cilia were also observed (**Fig. S5B**). At DIV90, we observed an initial but consistent S-opsin signal spanning the entire tissue, with some signal concentrated at the most apical side. Additionally, we noticed a distinct and well-defined Peanut agglutinin (PNA) staining in the emerging bud-like OS of the immature cones (**Fig. 3D-arrows**). When isolated at DIV60 and grown in suspension cultures, RS punches exhibit a natural tendency to fold inwards, forming organoid-like structures (**Fig. 3E**). At DIV150-DIV160, these were strongly positive for CRX, OTX2 and expressed apically PNA, RCVRN, S-opsin, and M/L-opsin (**Fig. 3F-G**), with relatively mature OSs resembling those found in the adult human retina (**Fig. 3H**). These results show that RPCs and PRs precursors present at DIV60 can develop into mature cones *ex vivo*.

### Generation of a porcine model of severe photoreceptor degeneration

Pig eye anatomy and dimensions closely resemble those of humans and include a visual streak enriched in cones, bearing some resemblance to the macula found in primates ^65^. We induced severe photoreceptor degeneration in this “pseudo-macula” of adult Yucatan minipigs through sub-retinal cobalt chloride injections (**Fig. 4A-B, and Video S1**)^23,66^. Utilizing a single apparatus, we conducted Optical Coherence Tomography (OCT) and multifocal Electroretinogram (mfERG) analyses under photopic conditions. These tests confirmed retinal lesions measuring 4-9 mm in diameter one month after treatment, with loss of visual function at the lesion site in correspondence with the ONL-free area (**Fig. 4C)**. Variations in lesion size and severity were primarily linked to solution reflux post-injection, affecting the final cobalt chloride volume in the retinal bleb (**Table S1**). One-month post-treatment, IF analyses confirmed complete loss of the ONL in the damaged area (**Fig. 4D-F**). While the RPE and INL generally remained intact, we observed more severe damage in some eyes, especially in the central portion of the lesion, which could lead to some RPE and INL loss (**Fig. 4D-F**). Staining for VSX2, which labels bipolar neurons in the mature retina, indicated a generally intact INL at the lesion site (**Fig. 4G**).

**Figure 4:**
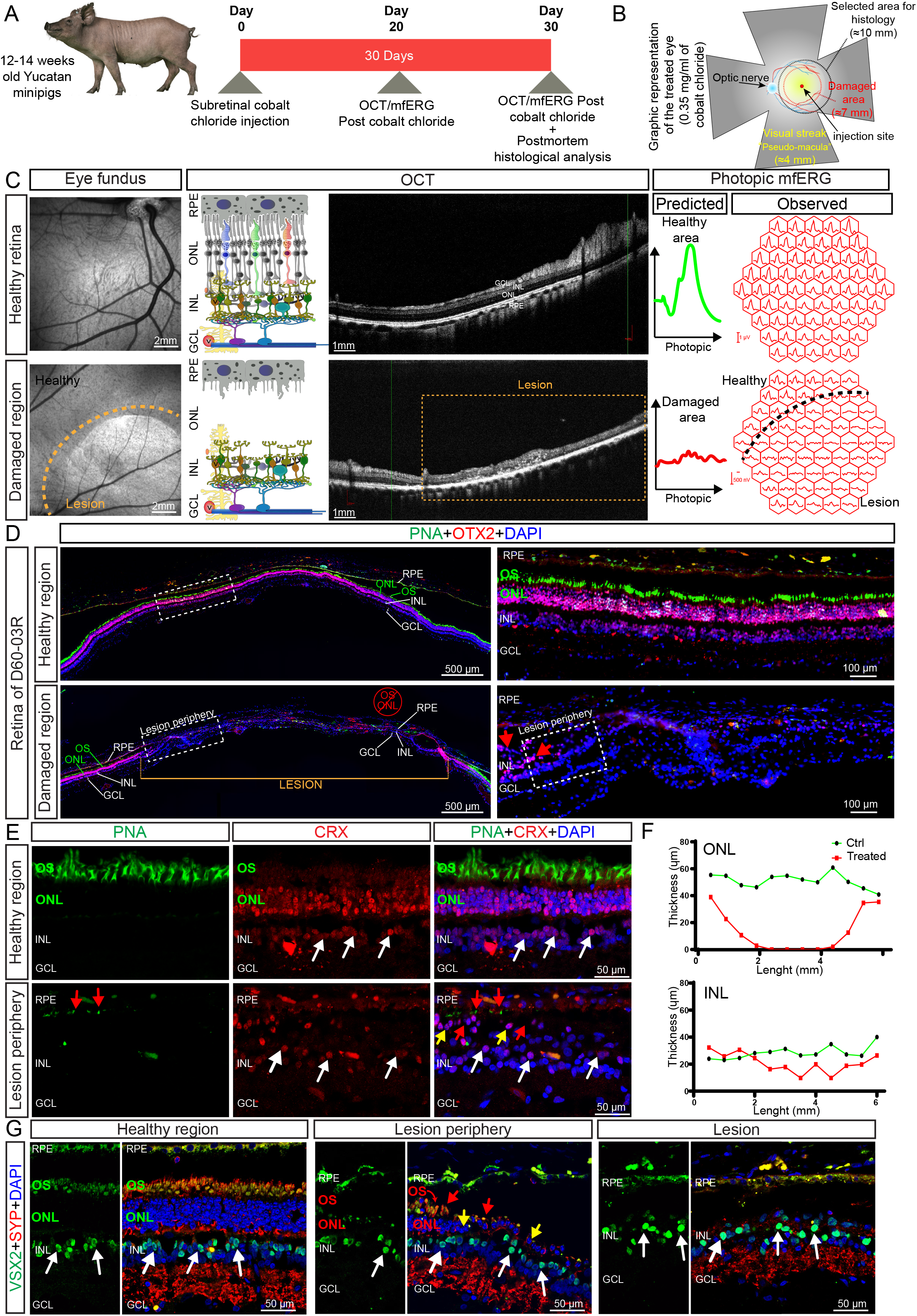
Generation of a minipig model of severe photoreceptor degeneration. A-Schematic representation of the study timeline. Yucatan minipigs (12-14 months old) were subjected to chemically induced macular degeneration using subretinal injection of 0.35 mg/ml cobalt chloride. 20 and 30 days post cobalt chloride injection, we conducted an OCT/ERG follow-up to assess the retinal damage. Animals were euthanized 30 days after treatment for histological analysis. B-Graphic representation of the anatomy of the minipig retina and the lesion upon treatment with 0.35 mg/ml of Cobalt Chloride. C-Comparison between eye fundus, OCT, and photopic mfERG from the healthy retina and the retina at D30 post cobalt chloride injection on the animal D60-03R pre-transplantation. In the OCT image and its graphical representation on its left, we can observe a complete loss of outer segments (OS) and Outer nuclear layer (ONL) in the lesion area. The yellow dotted box indicates the lesion site. On the right, mfERG recordings of the healthy and damaged regions and the predicted forms of the ERG signal in the two conditions. D-Representative immunofluorescence image of the healthy region and the lesion in the animal D60-03R. OTX2 labeling indicates photoreceptor and bipolar cell nuclei, while PNA stains cone OSs. A higher magnification of the periphery of the lesion site and the control area (white dotted rectangle in the left representation) is presented on the right. The yellow demarcated area is the lesion site. Red arrows indicate the few residual ONL nuclei at the margin of the lesion. The white dotted square in the right image indicates the area presented in E. E-Higher magnification of the periphery of the lesion site, where we can appreciate the rapid disappearance of the remaining photoreceptors nuclei (red arrows) and PNA signal (green arrows) from left to the right, in the D60-03R and the control eyes. White arrows indicate CRX-positive bipolar cells in the INL. F-Quantification of the thickness variation affecting the ONL and INL at the lesion site in the D60-03R eye and the control region. G-Representative immunofluorescence image of the healthy region, the lesion periphery, and the center of the lesion in the animal D60-03R. Bipolar cells nuclei are stained with VSX2, while photoreceptors’ OS and the inner and outer plexiform layers are stained with Synaptophysin (SYP). C, D, E, F Representative of n□=□20 □independent eyes. Scale bars are indicated in each figure.

### Grafted retinal tissue can integrate the minipig retina and generate a new ONL-like structure

To overcome a robust innate immune response previously observed in pilot experiments, minipigs were immunosuppressed using a multiple-drug regimen starting one week before surgery and maintained throughout post-transplantation (**Fig. 5A, and Material and Methods**). For transplantation, we generated a sub-retinal bleb of ∼5 mm diameter at the lesion site using a saline solution and subsequently injected one or multiple RS punches of 1-3 mm in diameter by shock waves using a custom-made injector (**Fig. 5B, Table S1, and Video S2**).

**Figure 5.**
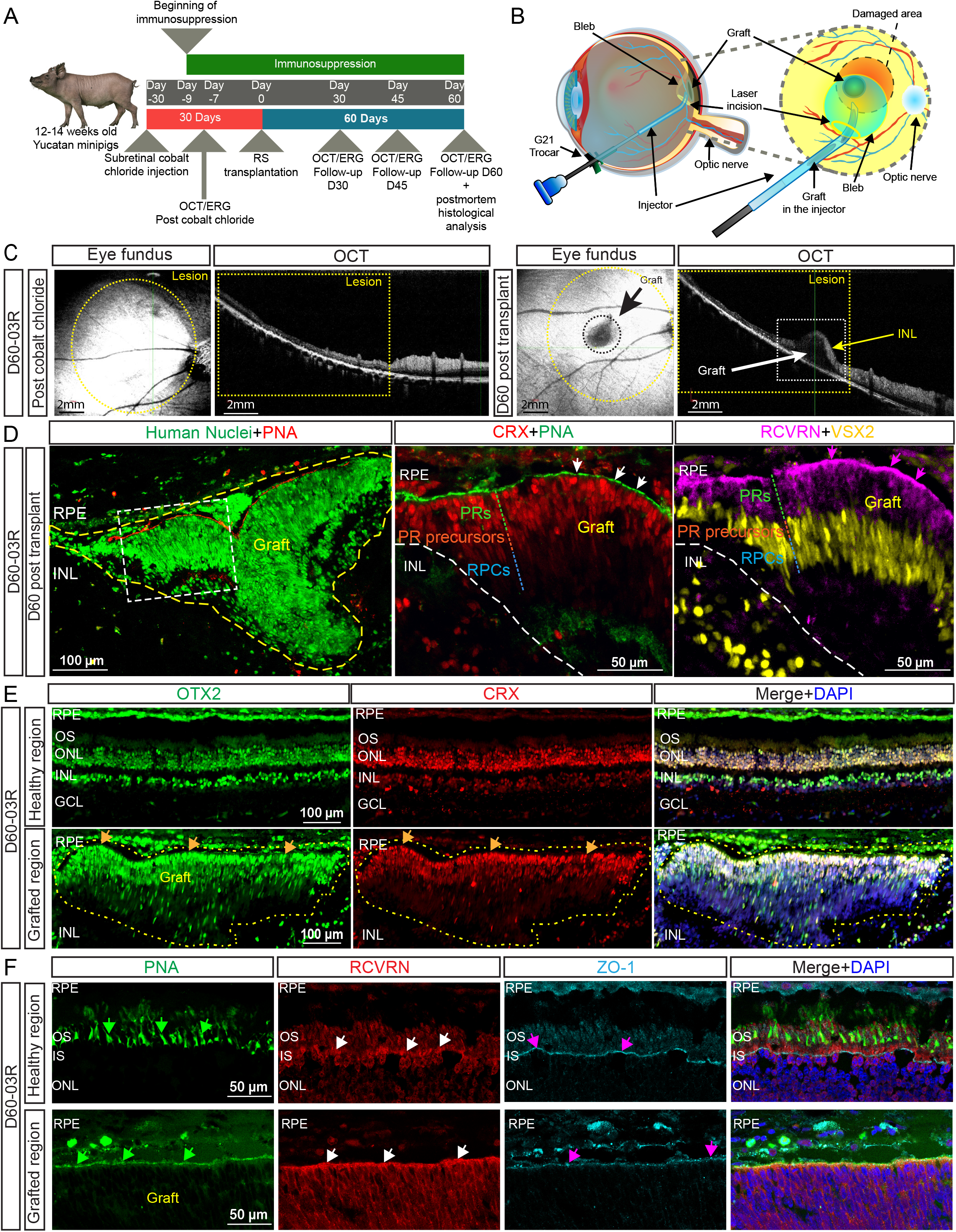
Grafted human retinal sheet can form a new ONL-like structure within the minipig retina. Schematic representation of the study timeline. Yucatan minipigs (12-14 months old) were subjected to chemically induced macular degeneration using subretinal injection of 0.35 mg/ml cobalt chloride. 23 days post cobalt chloride injection, we conducted an OCT/ERG follow-up to assess the retinal damage and select the animals suitable for transplantation. At D30 post cobalt chloride, we performed the transplantation of a 1,5-3 mm diameter punch of DIV60 retinal sheets using the WTC^Act^ or the clinical grade WTC^UDC^ cell lines. We conducted OCT and ERG follow-ups at D30, D45, and D60. The immunosuppressive regimen started from - D9 until the sacrifice (D60). B-Schematic representation of RS transplantation on the damaged area using a custom-made 20G soft-tip injector. C-OCT on animal D60-03R before the surgery and at D60 post-transplantation. The eye fundus shows that the graft is present within the lesion area (yellow circle), and the OCT shows a well-integrated graft (black dotted circle and white dotted box) with no ONL left in the damaged area (yellow dotted box) and an intact INL that covers approximately half of the graft (yellow arrow). D-On the left, the IF of the whole graft (yellow borders) was stained with human nuclei antibody to elicit its human origin and the PNA to show the orientation of the graft. On the right is a higher-magnification image of the graft integrated between the host RPE and INL. We can notice that the apical side of the graft is delimited with PNA and RCVRN. The CRX+ PRs reside on the apical side, and VSX2+ RPCs on the bottom, with a transition region that is CRX+VSX2+ identifying the immature PR precursors. E-Low magnification image of the graft 60 days post-transplantation. Note the strong expression of OTX2 and CRX within the flat portion of the graft (yellow borders), as well as its unique organization resembling a genuine embryonic ONL. Orange arrows indicate CRX+ OTX2+ double positive immature photoreceptors at the apical side of the graft. F-High magnification of the graft 60 days post-transplantation shows the junction between the graft and the endogenous retina. Note the clear demarcation borders delimited by PNA, RCVRN, and ZO-1. White arrows indicate apical RCVRN accumulation. The purple arrow indicates ZO-1 positive tight junctions. Green arrows indicate polarized PNA staining at the apical side. Scale bars are indicated in each figure.

At the time of surgery, the circular retinal lesion was visualized by fundus observation, and we could confirm the persistence of the graft at the lesion site by OCT analysis at day 30, 45, and 60 post-transplantations (**Figs. 5C, S6, and Table S1**). OCT and histological analyses revealed one or multiple issues with most grafts, including folding onto themselves (D60-03L, D60-09R, D60-09L), inverted apicobasal orientation (D60-04L), placement within severely damaged areas with affected host INL (D60-05R, D60-04L) or degeneration of the graft for its proximity to the incision site resulting in infiltration and partial destruction by macrophages (D60-07L). Another common issue was the positioning of the graft in the periphery or outside the fully damaged area (D60-01L, D60-04R), preventing proper interaction with the host retinal tissue and meaningful functional studies (**Fig. S6**). Histological analysis of the D60-03R eye however revealed the presence of a flat graft forming a relatively uniform retinal tissue within the host sub-retinal space and expressing the human-specific markers Human Nuclear Antigen (HuNu) and STEM-121 (**Figs. 5D and S7C**). We observed that two portions of this graft were unfolded and properly oriented with CRX and PNA-positive cells of the graft located apically and in close association with the host RPE (**Figs. 5D-inset and S7A**). While photoreceptor markers, such as CRX and RCVRN, were expressed apically in the entire and new ∼3-4 nuclei tick photoreceptor-like layer (**Fig. 5D)**, the formation of proper OSs was not observed, and only low levels of S-Opsin and M/L-Opsin were visible at the apical region of the graft (**Fig. S7D**). Within the graft, we noticed a complementary distribution of CRX and VSX2, with VSX2-positive retinal progenitors located inward of the graft and facing the endogenous INL (**Fig. 5D**). Some of these retinal progenitors exhibited low levels of Ki67 expression while testing negative for PCNA (**Fig. S7E**). This state is commonly observed during development in quiescent adult stem cells and progenitors, which remain in a non-proliferative resting state while remaining metabolically active and prepared to re-enter the cell cycle ^67–69^. Between these 2 populations, we observed the presence of presumptive photoreceptor precursors co-expressing CRX and VSX2 (**Fig. 5D-inset and S7A-B**). Notably, expression levels of CRX and OTX2 in this human iPSC-derived graft were similar to that of the adult pig’s endogenous photoreceptors (**Fig. 5E**). The adherents and tight junction protein ZO-1 demarcates the outer limiting membrane located between the photoreceptor’s cell body and the IS ^70–72^. Histological analyses revealed that the graft contained genuine but immature photoreceptors having a ZO-1, RCVRN, and PNA-positive apical edge in direct contact with the host RPE (**Fig. 6F**).

**Figure 6.**
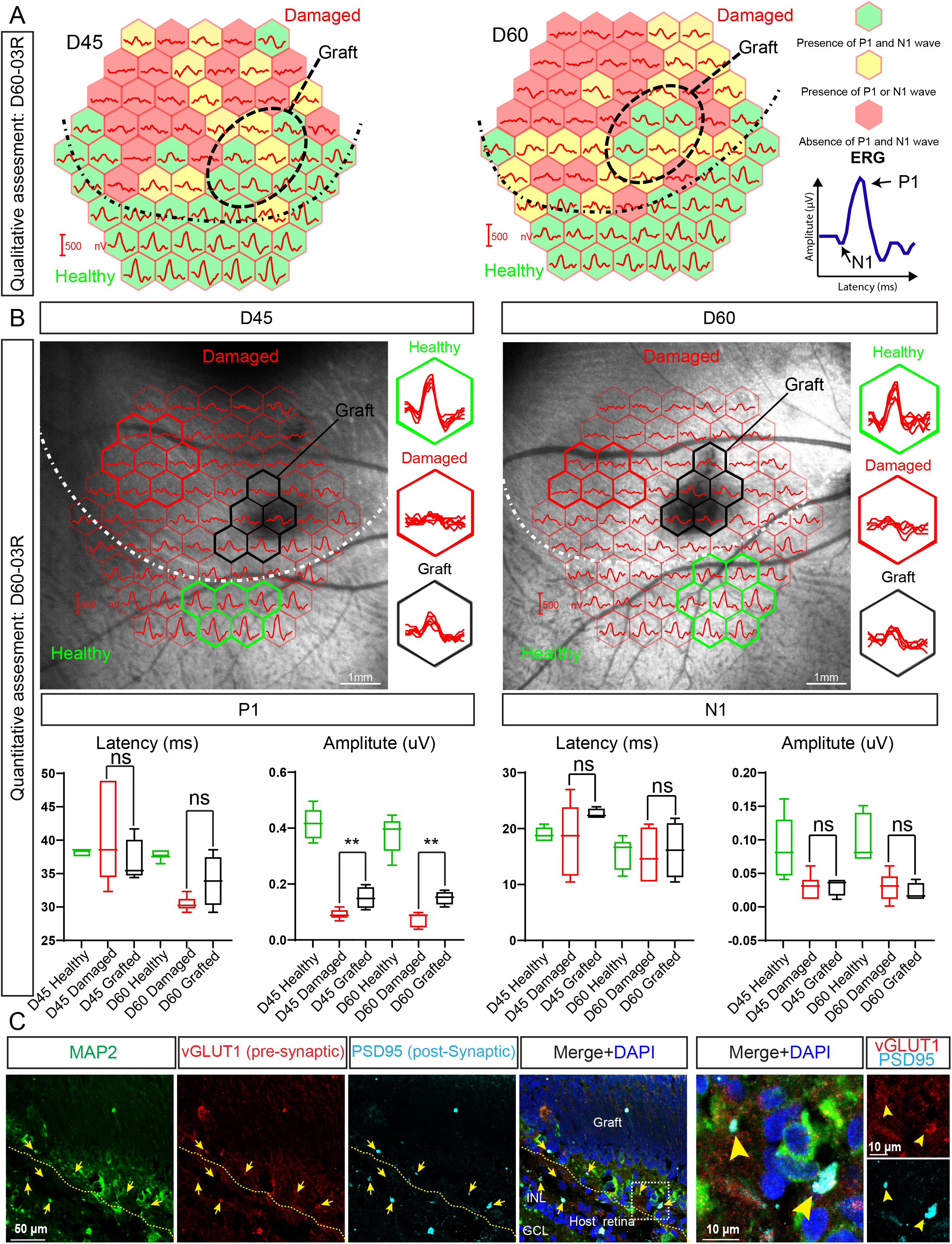
Evidence of functional integration of the graft within the host retina. A-A qualitative assessment of photopic mfERG on animal D60-03R at D45 and D60 post-transplantation. The signal in each hexagon is represented in the green, yellow, and red color codes. A dashed oval delimits the graft region. The dash and point line indicates the border between healthy and damaged areas.B-mfERG analysis on animal D60-03R at D45 and D60 post-transplantation. The eye fundus shows the graft in the black hexagons. The graft is present within the lesion area delimited by a white dashed line. The quantified regions are highlighted by a color-coded hexagon: red for the damaged region, green for the healthy region, and black for the graft. Quantified regions are superposed in the hexagons at the right of each fundus image to assess the generalized shape of the waves in different conditions. The graphs at the bottom show the quantifications of the latency and the amplitude of the P1 and N1 signals of highlighted regions.C-Confocal immunofluorescence images of the area between graft and host INL(delimited by the yellow dotted line) using pre- and post-synaptic markers. The arrows in yellow indicate the colocalization of pre- and post-synaptic markers vGLUT1, PSD95, and MAP2. On the right, a higher-magnification image of the area in the white dotted box. All values are means ± SEM. ∗p < 0.05, ∗∗p < 0.01 by Student’s unpaired t-test.Scale bars are indicated in each figure.

### Assessing graft function through mfERG recordings in photopic conditions reveals partial but significant rescue of P1 amplitude in the damaged area

To assess graft function, we conducted post-surgery follow-up analyses using multifocal electroretinogram (mfERG) recordings under photopic conditions to specifically measure the cone response, excluding the rod contribution. These assessments were attempted in all transplanted animals at days 30, 45, and 60 post-transplantations, except in cases where technical challenges, such as cataract formation or eye movement during analysis, prevented data collection (**Table S2**).

We generated a semi-quantitative binary (YES or NO) cumulative score sheet combining the above parameters: 1) quality of the retinal lesion (absence of the host ONL by OCT visualization and absence of mfERG response); 2) successful transplantation surgery; 3) graft localization within the lesion; 4) positive mfERG response of the retina at the graft position 60 days post-transplantation; 5) presence of an unfolded area within the graft (**Table S2**). From the initial number of WTC^UDC^ grafted animals (n = 10), 2 had cataracts, 2 had no graft within the lesion, and only 6 had a proper lesion (partial or complete) with a graft near or at the lesion site (**Tables S1 and S2**). After a more comprehensive postmortem immunofluorescence analysis, some of the grafts that initially exhibited functional activity in the mfERG analysis at the grafted site were found to have residual host ONL and had migrated to an area with partial or no damage at the border of the atrophic area. Consequently, samples such as the D60-04R, D60-07L, and D60-09R were excluded from the functional study as false positives (**Table S2, and Fig. S6**). From the initial number of WTC^Act^ grafted animals (n = 5), 2 had cataracts, and only 3 eyes were analyzed for OCT and ERG (**Tables S1 and S2**). Of all eyes (n = 30), only one grafted with the WTC^UDC^ cell line obtained a score of 10/10 (D60-03R), fulfilling all the requirements for potential functional integration into the host retina (**Table S2**). We then focused on the mfERG recordings at D45 and D60 from both the D60-03R and D60-03L eyes (**Figs. 6A and S8A**). The former had a well-polarized and flat graft, while the latter had a folded and improperly polarized graft (**Figs. 5D-E and S8C**). We performed a qualitative analysis to determine the presence or absence of N1 and P1 waves. N1 and P1 waves represent retinal responses to visual stimuli, with N1 indicating the initial negative deflection mainly associated with cone photoreceptor hyperpolarization. In contrast, P1 indicates the positive deflection associated with the depolarization of bipolar cells and inner retinal neurons ^73,74^. In the healthy control region of the host retina, both P1 and N1 waves were observed (green hexagons), while the damaged area often lacked both waves (red hexagons) or one of the two waves (yellow hexagons). Within the grafted region (highlighted by the dotted circle, consistently identified between days 45 and 60 using OCT and fundus imaging), traces consistently indicated the presence of N1 and P1 waves. (**Fig. 6A**). Next, we quantified the latency and the amplitude of waves between healthy, damaged, and grafted areas. We observed a partial but significant rescue of the P1 amplitude in the grafted area starting from day 45 and maintained till the endpoint at day 60 (**Fig. 6B**). This result is notable since the host photoreceptor ONL was completely ablated at this position before transplantation (**Figs. 5C and 4C-F**). Limited N1 response can be attributed to its intrinsic characteristics since it is much smaller than the P1 wave. When we conducted similar analyses on the D60-03L eye, which showed similar expression levels of photoreceptor markers like CRX, PNA, and RCVRN as the D60-03R eye but was completely folded and improperly polarized, no functional improvement was observed (**Figs. S8A-C**). These results suggest that the graft’s proper unfolding and orientation are crucial for visual function recovery.

### Evidence of synapse formation between grafted photoreceptors and host neurons

By IF, we observed a robust signal of the pre-synaptic marker synaptophysin within the basal portion of the graft in close proximity with the VSX2-positive bipolar neurons of the host (**Figs. S7B and S9A**). Consistent with the positive mfERG results in the D60-03R eye, we detected colocalization of the pre-synaptic and post-synaptic button markers vGLUT1 and PSD95, respectively, in the outer plexiform layer between the basal side of the graft and the host bipolar neurons (**Fig. 6C**). Similar observations were made when using the pre-synaptic markers vGLUT1, Synapsin, Synaptophysin, and CTBP2 (Ribeye) in combination with the post-synaptic markers PKC-alpha, MAP2, and PSD95 **(Figs. 6C and S9A-D**). Collectively, these results indicate the potential formation of synaptic connections between the grafted photoreceptors and the host bipolar neurons.

### Enhanced retinal graft survival and reduced immune reaction with the WTC^UDC^ iPSC line

Recruitment of microglial cells (IBA1+) was frequently observed near the injection site and around the grafted area (**Fig. S10A-C**)^75^. The innate immune response observed in eyes grafted with punches from the WTC^Act^ cell line was generally more robust compared to those from the WTC^UDC^ cell line, with significant infiltration of macrophages and IBA1+ activated microglia within the graft (**Fig. S10A-D**). Despite immunosuppression, we observed the destruction of 4 out of 7 grafts generated with the WTC^Act^ cell line (**Fig. S10D, and Table S2**). Longitudinal OCT analyses of eyes grafted with punches from the WTC^Act^ cell line also revealed thinning of the graft over time. We noticed a significantly lower immune reaction in eyes grafted with punches from the WTC^UDC^ cell line (**Fig. S10A-D**). Although there were few IBA1/CD4-double positive macrophages (anti-inflammatory M2 healing macrophages), acquired immune response (CD4 and CD8) was about absent 60 days post-transplantation in WTC^UDC^-grafted eyes (**Fig. S10A-D)**^76,77^.

## DISCUSSION

In this study, we have confirmed the capacity of 3 isogenic human iPSC lines to differentiate into RSs containing about all cell types present in the human eye, including RPCs and cone precursor cells. RSs were responsive to bright light stimulation *in vitro*, as shown using the cGMP degradation assay. Developmental kinetic analysis using scRNA-seq revealed that while present at DIV5, pluripotent stem cells were absent from DIV30 and DIV60 RSs, as further confirmed by the absence of teratoma formation in transplanted NOD/SCID mice. RS punches were found to be highly enriched for RPCs and photoreceptor precursors minus horizontal, amacrine, RPE, lens, and corneal cells, making them more suitable for photoreceptor transplantation therapy. RS punches in suspension cultures could differentiate into mature S- and M/L-cones *ex vivo*, while sub-retinal transplantation in minipigs revealed their capacity for integration and survival within the damaged host retina.

When compared to the developing human retina, DIV60 RS punches most resemble D-67-D80 human fovea based on the RNA-seq and histological data ^29^. In the human fovea, OTX2 labels the one nucleus thick ONL and the much larger INL that also expresses VSX2. A 4-5 nuclei thick ganglion cell layer is also present and that is negative for both markers ^29^. While a near identical OTX2+ ONL is also present in the RS punches, the presumptive INL only partially co-expresses OTX2 and VSX2, and the ganglion cell layer is indistinguishable with only ∼2% of RGCs being present (**Figs. 2A’ and 3B**). DIV60 RS punches also share a comparable developmental state with freshly dissected human fetal retina sheets used in subretinal transplantation studies ^78–80^. These studies have consistently demonstrated promising outcomes in both animal models and human clinical trials, characterized by the preservation and restoration of visual responses ^78–80^. Nevertheless, they have not been adopted as standard clinical practice due to ethical constraints associated with the utilization of human fetal tissue. The pivotal role of synaptic connections between the transplant and host in achieving this visual improvement has been well-documented. Notably, phase II trials in patients with RP and ARMD have demonstrated substantial enhancements in visual acuity following transplants of fetal retinas along with their retinal pigment epithelium (RPE)^79,81,82^. Intriguingly, it has been observed that immunosuppression may not be requisite for fetal retina transplantations if the intact retinal barrier is preserved during surgery^79,81,82^. Our iPSC-derived RSs, while possessing a similar cellular composition to the fetal retina, present a distinct advantage for macular transplantation because apparently lack rods, bipolar, amacrine, and horizontal cells (**Fig. 2D**). Additionally, the ease of handling RSs offers the potential for integration with RPE monolayers, providing a promising solution for cases involving concurrent damage to both photoreceptor and RPE cells.

As suggested by previous studies, fully mature cells often display limited integration capacity within host tissue, while excessively immature progenitors may fail to properly mature into a polarized and mature layer, potentially resulting in *in vivo* cell proliferation^83,84^. Herein, we chose to employ DIV60 RS punches for retinal transplantation. These cells have predominantly lost their proliferative traits and consist of progenitors and precursors largely poised for photoreceptor cell fate. Interestingly, unlike observations from other studies^16^, we did not observe enhanced graft maturation due to donor-host interaction *in vivo*; instead, grafts followed a developmental pace relatively similar to the *ex vivo* cultures. The current experimental approach involves a xenogeneic and heterochronic paradigm, as human embryonic tissue is transplanted into the retina of adult minipigs. Achieving full cone differentiation and the development of completely mature outer segments (OS) in the transplanted animals may necessitate an additional six months ^85–87^. By extrapolation, prospective patients treated with this regenerative medicine approach may require up to 1 year before observing optimal therapeutic effects.

We built a “performance scoring sheet” of all grafting experiments, which revealed a strong correlation between the presence of a polarized photoreceptor sheet *in situ* and a functional response at the grafted site. The correlation between the structure of the graft and its function helped us define key elements for its successful engraftment. Hence, proper differentiation, polarization, and functionality of the newly introduced photoreceptor layer critically depend on the uniform distribution of the graft as an unfolded retinal tissue. Graft’s folding resulted in the formation of rosettes-like structures, similar to what was reported from other groups using dissociated photoreceptors and retinal organoids ^15–17,26^. Growing RSs on a semi-solid biodegradable matrix or a flexible solid support may prevent graft folding during transplantation. The use of matrix or similar supports has proven effective in transplanting pluripotent stem cell-derived RPE monolayers^88–90^. Coupled with the utilization of more advanced delivery devices, this approach has the potential to significantly increase transplant success rates ^91^.

Although clinical-grade autologous iPSC lines can be created for each patient, the associated expenses and time requirements are considerable. Additionally, patients with inherited retinal degenerative diseases will require genetic correction of the identified mutation before the development of autologous iPSC lines. Consequently, the availability of hypoimmunogenic iPSC lines created worldwide offers promising possibilities for treating patients with retinal degeneration, circumventing the need for autologous iPSC and immune suppression^92–95^. This work paves the way for future long-term functional investigations utilizing RS grafts and holds great promise for the treatment of macular degeneration and retinal dystrophies.

## Supporting information

Video 1-Cobalt Chloride subretinal injection

Video 2-Retinal Sheet Transplantation

## LIMITATIONS OF THE STUDY

The surgical procedure to ensure correct graft localization, apicobasal orientation, and unfolding needs to be improved. The small number of surgeries answering all of these requirements represents the main limitations of this study.

## ACKNOWLEDGMENTS

This work was supported by grants from the Stem Cell Network-Canada (FY19/DT9 and FY20/FBP-1), donations from the Felicia and Arnold Aaron Foundation, and partnerships with StemAxon and Healios KK corporations. The initiation of this project was supported by a historical donation from the Turmel Family Foundation for macular degeneration research. Thanks to Bausch Health Canada and Joe Matar for essential technical support in this project, and to Drs Flavio Rezende and Cynthia Quian for their initial support and implication in this study.

## DATA AVAILABILITY STATEMENT

Raw data, cell lines, and reagents are available upon request.

## CODE AVAILABILITY STATEMENT

Accession code for scRNA-seq data is GSE228436

## AUTHOR CONTRIBUTIONS

Conceived and designed: G.B., A.B., K.M., J-F.B., M.G., and R.H.

Performed the experiments: G.B., A.B., K.M., A.K., A.P., and R.H.

Analyzed the data: G.B., A.B., R.H., A.P., and K.M.

Wrote the paper: G.B., A.B., K.M., and R.H.

## MATERIAL AND METHODS

### Cell cultures

The isogenic WTC^Act^ and WTC^Tub^ (Allen Institute) and WTC^UDC^ iPSC lines, and the iPSC#1 and iPSC#2 iPSC lines, and the H9 (WiCell) hESC line were dissociated using ReLeSR™ (Stem cell technologies, Cat#05872) and platted on growth factor reduced iMatrix in StemFlex cell media (Gibco # A3349401), supplemented with ROCK inhibitor (Y-27632;10_M, Cayman Chemical #10005583). WTC^UDC^ is a hypoimmunogenic line with inactive α-chain HLA Class 1a and HLA Class II genes expressing the exogenous α-chain HLA Class 1b, PD-L1, and PD-L2 genes (WO2021241658A1). The iPSC#1 and iPSC#2 lines were generated by reprogramming normal human fibroblasts using non-integrative plasmids expressing the Yamanaka factors and a small hairpin-RNA against p53. Upon confluency, cells were differentiated with CI media: DMEM-F12 medium (Invitrogen) containing 1% N2, 2% B27, 10 ng/ml IGF1 (PeproTech, Cat#100-11), and 5 ng/ml bFGF (PeproTech, Cat#AF-100-18B), Heparin (Sigma, Cat#H3149), and 30 ng/ml COCO (R&D System, Cat#3047-CC-050) ^27^. DIV60 punches were cultured in CI medium as suspension cultures in ultra-low adherence plates (Corning, CLS3473).

### RNA-seq analysis

Total RNA from two independent biological replicates was extracted using the standard procedure of Qiagen columns and assayed for RNA integrity. cDNA was prepared according to the manufacturer’s instructions (NEB library) and sequenced using the Illumina platform. Base-calling and feature count were done using Illumina software. For differential expression analysis, Dseq2 ^96^ was used on the R program. The samples were analyzed along a public dataset of bulk RNA at various stages of the human fetal retina. A heatmap was plotted with the photoreceptor genes to match the maturity of our retinal sheets with the developmental stage of the fetal retina.

### Single-cell RNA-seq analysis

Cells were dissociated and prepared with the standard procedure of 10X Single-cell gene expression kit aiming for the collection of 10,000 cells. The resulting library was sequenced with the Illumina platform. Base-calling and feature count were done using Illumina software. Read count was performed with Cell Ranger software 3.0.0^97^ from 10X to analyze and map the reads on the HG38 genome. The counts of this experiment were aggregated with the analyzed data from the publicly available Descartes dataset of the eye using the Seurat package ^98^ The Descartes dataset is a published dataset of the human fetal retina ^99^. Using anchor points between these two datasets, we plotted them on the same 2D space using UMAP^100^ distribution to understand our cells’ identity. Monocle3 ^101^ was used to calculate the pseudo-time ^102^ between the DIV5, DIV30, and DIV60 RSs. In summary, the three datasets were combined using Monocle3, then a UMAP ^100^ was generated from the combined dataset. On those data, Monocle3 was able to learn a differentiation graph that would allow us to plot the pseudo-time. Loupe 4.1.0 ^103^ was instrumental in visualizing the data and performing an unbiased ontology analysis to identify the cell lineage present in our cultures. In summary, cells were clustered without bias using the K-mean strategy. The differentially expressed genes in each group identified by the cell ranger were compared to the literature and their ontology origins to identify the corresponding group.

### Minipig model of retinal degeneration and subretinal transplantation

See the Supplementary Material and Methods document.

### Immunosuppression regimen

The immunosuppressants to be used for all groups are the following:

-Starting 9 days before Transplant (-D9) and continuing until euthanasia:

∘ Doxycycline 10mg/kg SID PO
∘ Methylprednisolone 5mg/kg IM on -D9 only, followed by administration of Prednisone 5mg/kg SID PO starting on -D8
∘ Tacrolimus 0.5mg SID PO
∘ Triamcinolone (Kenalog) 40mg/mL 2.8mg/eye IVT on D0

### OCT and mfERG

Animals received Mydfrin 2.5% TP and Cyclogil 1% TP approximately 60 minutes, 45 minutes, and 15 minutes before the surgery to induce mydriasis. Drops of Mydriacyl 1% were also given prior to the imaging procedures to optimize mydriasis. Optixcare eye lube will be applied to keep the eyes lubricated during the imaging procedures. The animals were anesthetized by administration of butorphanol (0.20 mg/kg), ketamine (10 mg/kg), and dexmedetomidine (0.04 mg/kg) given IM. Upon induction of anesthesia, animals will be intubated and supported with mechanical ventilation. A corneal instillation of Lidocaine hydrochloride 2% was given in each eye. A Jet-Electrode (Roland Consult) was placed on the eye with an appropriate amount of Optixcare eye lube. OCT was performed using the RETImap animal system (Roland Consult). The region of interest (ROI) was placed in the center and an averaged single scan across ROI was performed. mfERG was recorded following the OCT, using the RETImap system (Roland Consult). Briefly, a Jet-Electrode was placed on the eye, reference, and ground electrode (Genuine Grass Platinum Subdermal Needle Electrode (Natus Manufacturing Limited) were placed respectively near the ipsilateral outer canthus and on the chin. A minimum of 10 cycles were used for each recording. A low bandpass of 10hz and a high bandpass of 300 Hz were used. The mfERG waveform includes an initial depolarization (N1), followed by a repolarization (P1). The N1 wave reflects the initial negative deflection corresponding to cone photoreceptor cell activity. This wave component directly measures the cone’s function. On the other hand, the P1 wave is the amplified response generated by bipolar and amacrine cells in response to the cone’s activity (N1), thus reflecting phototransduction by the whole cone system. Interestingly, while N1 can exist in the absence of P1, as it occurs when bipolar cells are destroyed or nonfunctional and photoreceptors are preserved, P1 cannot exist in the absence of N1. Due to its low amplitude, N1 is more challenging to detect and is lost faster during photoreceptor degeneration^73,74,104^. For qualitative assessment of the mfERG, all the traces were separated and anonymized and then scored by 5 different persons that had the same guidelines: if the trace had no N1 or P1 waves score the score is 0, if the trace had either N1 or P1 wave the score of that trace is 1, and if the trace had both N1 and P1 waves the score of the said trace is 2. The judges were also given a good standard trace and a standard flat trace for comparison. The results were then merged and averaged and then traced back to the appropriate hexagon on the reading. For quantitative assessment of the mfERG, we quantified the latency and the amplitude of the N1 and P1 waves of a subset of hexagons. For the healthy and damaged area, a central hexagon was chosen that had its border well defined in the corresponding area (damaged or healthy); we quantified this hexagon and all its surrounding hexagons if they were not on the border between the healthy and damaged area. For the graft, we quantified the hexagons that were visibly overlapping the graft.

### Preparation of the ocular tissues

Thirty days after surgery, Yucatan minipigs were euthanized by inducing deep anesthesia followed by a lethal injection of pentobarbital (Euthanyl Forte 540 mg/mL 10 mL/50kg) as a rapid bolus. Both eyes from each treated animal were enucleated, incisions were made through the pars plana, and the globes were immersed in 4% paraformaldehyde in PBS (Sigma-Aldrich Corp.) on ice. The anterior chambers were then removed and the vitreous removed, and the eyecups were fixed for another 10–30 min in the same fixative depending on how much vitreous was left. After three washes of 10 min each in 1· PBS, the minipig eyecups were cryoprotected in sucrose solutions prepared in PBS, pH 7.8 (15% sucrose for 1–2 h until sinking, then 30% sucrose for an hour, and finally 50% sucrose O/N). The minipig eyecups were then snap-frozen in optimal cutting temperature embedding material (Neg50 Frozen Section Media, ThermoFisher) in a beaker filled with dry ice. They were then stored at -80°C until sectioned. The cryostat (Leica) was used to produce 10 μm serial sections of minipig eyes.

### Immunofluorescence

Retinal cryosections are permeabilized with Triton X-100 for 10 min. Unspecific antigen blocking was performed using 2% BSA in PBST for 30 min. Slides were incubated with the primary antibody overnight at 4°C in a humidified chamber (see dilutions in Table). Secondary antibodies were incubated for 1h at room temperature. After incubation with the secondary antibody, slides were washed, counterstained with DAPI and mounted using VECTASHIELD® Antifade Mounting Medium (VECTASHIELD, H-1000-10) and No. 1.5H coverslips. Pictures were taken using a confocal microscopy system (Olympus). Confocal microscopy analyses were performed using 60x objectives with an IX81 confocal microscope (Olympus, Richmond Hill, Canada), and images were obtained with Fluoview software version 3.1 (Olympus).

### cGMP assay

Phototransduction activity was assessed by measuring the light-induced hydrolysis of endogenous cyclic (c) GMP with an enzyme immunoassay kit (Biotrack EIA system) according to the manufacturer’s instructions (Amersham Bioscience GE Healthcare). Undifferentiated WTC^Act^ or WTC^UDC^ iPSC and cells differentiated into RS were kept in the dark. The cells with the “light condition” were exposed to 5 min of bright light before the analysis. The PDE inhibitor IBMX (3-isobutyl-1-methylxanthine, Sigma) was added (1 mM) 72 h before the determination of cGMP levels.

### List of antibodies

A list of antibodies can be found in the Supplementary Material and Methods document.

