## Supplementary materials and S.figures legends for "Sub-retinal transplantation of human iPSC-derived retinal sheets: A promising approach for the treatment of macular degeneration"

### SUPPLEMENTARY FIGURE LEGENDS

- **Figure S1: Molecular characterization of iPSC-derived retinal sheets**

A- Overview strategy of the study with key deliverables.

B- Flow cytometry analysis of $WTC^{UDC}$ clinical-grade retinal sheet at DIV60 using CRX marker (AF488). More than 65% of cells are CRX-positive, divided into two populations expressing low (53.6%) or high (11.9%) levels of CRX.

C- Heatmap of bulk RNAseq comparing retinal sheets differentiated for 60 days from the $WTC^{UDC}$, two different control iPSC, and one hESC (H9) with the bulk RNA seq of the human fetal retinal at various stages of development. The expression pattern of our 4 differentiations resembles the pattern seen on days 67-94 in the human fetal retina.

D- cGMP analysis of iPSC and retinal sheets at day 60 of differentiation for the $WTC^{Act}$ and $WTC^{UDC}$ lines. On the right, the PDE inhibitor-treated (1mM) control for the $WTC^{UDC}$ cell line. All values are means ± SEM. *$p < 0.05$, **$p < 0.01$, by Student's unpaired t-test.

B, C, D Representative of n = 3 independent experiments.

- **Figure S2: Gene expression profiling of retinal sheets at different stages of differentiation by scRNAseq analysis.**

A- Feature plot representation of scRNAseq data showing UMAP graphs of the $WTC^{UDC}$ line at DIV5, DIV30, or DIV60 of differentiation for genes highlighting retinal fate: SIX6, RAX, and VSX2 (CHX10). Each dot on the plot represents a single cell and its corresponding transcriptional activity levels.

B- Feature plot representation of scRNAseq data showing UMAP graphs of the $WTC^{UDC}$ line at DIV5, DIV30, or DIV60 of differentiation for genes highlighting optic stalk fate: PAX2, VAX1, and NKX6.2. Each dot on the plot represents a single cell and its corresponding transcriptional activity levels.

C- Feature plot representation of scRNAseq data showing UMAP graphs of the $WTC^{UDC}$ line at DIV5, DIV30, or DIV60 of differentiation for genes highlighting photoreceptor fate: RXRG, RCVRN, PDC, PDE6H, ARR3, and GNGT1-2. Each dot on the plot represents a single cell and its corresponding transcriptional activity levels.

D- Heatmap of the most dysregulated genes between the cell types at DIV5. Each row represents a cell type, and each column represents a distinct gene. These genes can be grouped according to the cell type in which they are mostly expressed, as shown with the color-coded bar above the heatmap. The colors follow the same code as in Fig1-F. We highlighted some genes that would help to identify and confirm the group identity.

E- Heatmap of the most dysregulated genes between the cell types at DIV30. Each row represents a cell type, and each column represents a distinct gene. These genes can be grouped according to the cell type in which they are mostly expressed, as shown with the color-coded bar above the heatmap. The colors follow the same code as in Fig1-F.

. **Figure S3: QC and safety assessment or iPSC-derived retinal sheets**

A- Feature plot representation of scRNAseq data showing UMAP graphs of the $WTC^{UDC}$ line at DIV5, DIV30, or DIV60 of differentiation for genes highlighting pluripotency markers POU5F1, NANOG, ZFP42, MYC, and KLF4. Each dot on the plot represents a single cell and its corresponding transcriptional activity levels.

B- UMAP graphs of the $WTC^{UDC}$ line at DIV5, DIV30, or DIV60 of differentiation highlighting in red the cells that are simultaneously positive for POU5F1 and: NANOG, ZFP42, MYC, or KLF4. The number of positive cells for each combination was counted and reported in the graph, along with the corresponding percentage.

C- Summary of the tumorigenicity assay in NOD/SCID mice using hESC and iPSC-derived retinal sheets differentiated for 60 days to assess the safety of our product. Undifferentiated hESC and iPSC are used as positive control.

. **Figure S4: Analysis of retinal sheets: biopsy punch vs. bulk**

A-E- Feature plot representation of scRNAseq data showing UMAP graphs of the WTCUDC line at DIV60 of differentiation for the bulk population and the area selected by punch biopsy. Highlighted genes are representative of: A) RPCs fate, B) late RPCs, C) RGC, D) retinal neuronal fate, and E) optic stalk. Each point represents the transcription levels of the gene in a single cell. These graphs show the preservation of these cell populations in the punch. A circle color coded with the cell type color highlights the position of these cell populations.

- **Figure S5:**

A- Confocal IF images of cryosections of retinal sheet punch at DIV60 generated from the WTC$^{UDC}$ GMP-grade cell line using specific photoreceptor and progenitor markers (CRX, OTX2, PNA, RCVRN, VSX2). Green arrows indicate the initial formation of a rudimentary OS (PNA+). White arrows indicate cone precursors CRX+$^{high}$ and OTX2+$^{high}$. Cyan arrows indicate RPCs (CHX10+, CRX+$^{low}$). Yellow arrows show apical accumulation of Recoverin (RCVRN) in photoreceptors precursors.

B- The figure shows a retinal sheet punch visualized using transmission electron microscopy (TEM). The white arrows indicate the presence of tight junctions between adjacent cells, while the black arrows indicate the budding of the IS of the immature photoreceptor cells. Asterisks indicate the location of basal bodies. M represents mitochondria, and N refers to the nucleus. Additionally, connecting cilia (CC) and basal bodies (BB) are shown in a transverse cross-section.

- **Figure S6: Overview of the transplantation results that highlight problematics encountered after transplantation.**

Summary results of the transplanted animals with the WTC$^{UDC}$ to highlight problematics incurred during and post-surgery. Eye fundus, OCT, photopic mfERG, and IF analysis on D60-01L, D60-03L, D60-04R, D60-04L, D60-05R, D60-07L, D60-09R and D60-09L at D60 post-transplantation. In the IF HuNu is used to stain the graft, CRX and PNA are used to mark the graft and the host retina, and S-Opsin is used to show the intact OS of the host retina over the graft in the eye D60-01L. Scale bars are indicated in each figure.

- **Figure S7: Immunofluorescence analysis of the graft in the D60-03R eye reveals a polarized and multi-layered transplanted retinal tissue**

A- IF of the graft at low magnification for PNA and CRX. The yellow dotted line delimits the graft. The white box shows the location of the islet shown on the right. The white arrows indicate the apical PNA staining above the CRX-positive nuclei.

B- IF of the graft at low magnification for Synaptophysin and VSX2. The white arrows indicate apical accumulation of synaptophysin, blue arrows indicate VSX2+ nuclei.

C- IF of the graft at low magnification for STEM121(human-specific marker) and RCVRN. The white arrows indicate apical accumulation of RCVRN in Stem121+ cells.

D- IF of the graft at low magnification for S- and M/L opsins (human-specific marker). The white arrows indicate apical accumulation of S- and M/L opsins.

E- IF of the graft at low magnification for PNA and Ki67 (proliferation markers). The white arrows indicate Ki67+ nuclei while no cells are positive for PCNA.

The yellow dotted line delimits the graft. The white box shows the location of the islet shown on the right. PRs, PR precursors, and RPC layers are indicated in the image with dotted lines. Scale bars are indicated in each figure.

- **Figure S8: Graft's polarization and unfolding are critical for its functionality**

A- mfERG analysis on animal D60-03L where the patch was folded and not properly oriented at D45 and D60 post-transplantation. A white dashed line delimits the lesion area in the eye fundus. Graft is included in the black hexagons. The quantified regions are highlighted by a color-coded hexagon: red for the damaged region, green for the healthy region, and black for the graft. Under the eye fundus, we can see a visual superposition of these quantified regions to assess the generalized shape of the signal in each condition.

B- Graphs showing the quantifications latency and amplitude of the P1 and N1 waves of the regions indicated in A.

C- Eye fundus, OCT, and IF analysis on the D60-03L animal showing the lack of a flat contact area between the graft and the host retina.

Scale bars are indicated in each figure.

- **Figure S9: Evidence of synapses formation between the graft and the host retina**

A-D- High-magnification confocal images of the synaptic markers (CtBP2, Synaptophysin, Synapsin, vGLUT, PKCα) focusing on the border between the graft and the INL of the host retina. STEM121 is used as a human-specific cytoplasmic marker. VSX2 stains the RPC population in the graft and bipolar cell nuclei of the host retina. Here we can notice the different expression patterns of VSX2 and the different shapes of the nuclei of the 2 populations. The yellow arrows indicate the putative synaptic connections shown by the colocalization of synaptic markers. White arrows indicate a STEM121 cytoplasmic extension of the graft penetrating the host INL. The dotted line represents the border between the graft and the host INL. The yellow dotted box shows the location of the islet shown on the right.

Scale bars are indicated in each figure.

- **Figure S10: Evidence of reduced immune response in grafts produced from the hypoimmunogenic iPSC line**

A- Immunofluorescence analysis of the innate immune response (IBA1) and the adaptive immune response (CD4/CD8) of the control retina and the grafted area 60 days after transplantation of $WTC^{UDC}$ and the $WTC^{Act}$ cell line. The yellow dotted line delimits the graft; the yellow arrows indicate IBA-1-positive activated microglia/macrophages. White arrows indicate double-positive IBA1 and CD4/CD8 cells.

B- Immunofluorescence analysis to assess cell proliferation in the grafted area with the $WTC^{UDC}$ and the $WTC^{Act}$ cell lines 60 days after transplantation. The ungrafted eye is used as a control. White arrows indicate PCNA-positive nuclei. Yellow arrows indicate KI67-positive nuclei.

C- Immunofluorescence analysis was conducted to assess the innate immune response (IBA1+) at the graft level and at the incision site generated during the surgery to introduce the injector under the retina. Remarkably, immune cell recruitment was found to be higher at the incision site than at the graft, indicating that it is primarily due to surgical trauma rather than the nature of the transplanted cells. The graft is delineated by the yellow line, with IBA1-positive cells denoted by yellow arrows and HuNu-positive cells marked by blue arrows.

D- Summary of the quantification of the proliferation, innate immune response (IBA1+), and adaptive immune response (CD4/CD8+) at the injection site and at the graft in all transplanted animals. Note the reduced immunogenicity of the $WTC^{UDC}$ compared to the $WTC^{Act}$ line.

Scale bars are indicated in each figure.

- **Video S1: Cobalt chloride subretinal injection.**

- **Video S2: Transplantation or RS punch in the subretinal space.**

  a) Injection of saline to generate the sub-retinal bleb.
  b) Injections of the 3 mm diameter grafts in the bleb.
  c) Yellow arrows indicate the 2 grafts inserted in the subretinal space.

### SUPPLEMENTARY MATERIAL AND METHODS

#### Minipig model of retinal degeneration

Sub-retinal injections of cobalt chloride hexahydrate (0.35 mg/ml in saline solution) were performed on 12-14 months old males Yucatan minipigs (n = 16). Animals were injected with 40µl of cobalt chloride following subretinal bleb formation on both eyes. Four weeks later, animals were examined by mfERG and OCT (Roland Consult).

#### Subretinal transplantation

All animal procedures were approved by the Institutional Animal Care and Use Committee of the Hôpital Maisonneuve-Rosemont Research Center and conducted in accordance with the Canadian Council on Animal Care (CCAC). All retinal surgeries were performed by a boarded ophthalmologist and vitreo-retinal surgeon. Minipigs were anesthetized with isoflurane and placed in ventral recumbency. The eye was positioned in primary gaze and aseptically prepared for a routine 3-port pars plana 25-gauge vitrectomy (BAUSCH and LOMB). A light pipe is introduced into the eye to allow direct visualization of the whole procedure under the retinal surgery microscope (Leica). A core vitrectomy using Stellaris PC (BL1433D, BAUSCH, and LOMB) and detachment of the posterior vitreous face over the region of planned implantation were performed. A subretinal injection of BSS was performed using a DeJuan/Awh 41-gauge subretinal injection cannula (BAUSCH and LOMB) to induce the formation of a bleb and a retinal detachment. For cobalt chloride and dissociated cells treatments, 40 ul of solution or cell suspension were injected using a DeJuan/Awh 41-gauge subretinal injection cannula without performing a vitrectomy and using a Hamilton syringe. For transplantation of the 3 mm biopsy punches (i.e., grafts), a 20G trocar was inserted to accommodate the entry of the injector, and a retinotomy was performed into the detached retina with retinal scissors to allow entry of the injector into the subretinal space. Grafts were kept at 37°C in a 5% $CO_2$-saturated tissue culture incubator in the culture medium until 10 min before implantation. Grafts in 150 µl 0.9% NaCl were loaded in the injector and injected into the subretinal space under direct visualization. Sclerotomies were closed using a 7–0 Coated Vicryl-Rapide suture (Ethicon, Inc). Laser catheterization of the retinal incision was made using Stellaris PC. Triamcinolone (3 mg/eye) was injected into the vitreous at the end of the procedure.

### List of antibodies

| Antibody | Catalog number | Company | IF dilution |
|---|---|---|---|
| CRX (CORD2 antibody) | GTX124188 | Genetex | 1:300 |
| S-OPSIN | AB5407 | Millipore | 1:200 |
| M-OPSIN | AB5405 | Millipore | 1:200 |
| Recoverin | AB5585-I | Millipore | 1:250 |
| PNA Rhodamine | VECTRL1072 | Vector Labs | 1:250 |
| PNA Fluorescein | VECTFL10715 | Vector Labs | 1:250 |
| IBA-1 | 019-19741 | Fujifilm WAKO | 1:300 |
| CHX10 | SC-21690 | Santa Cruz | 1:250 |
| PKC-a | SC-8393 | Santa Cruz | 1:250 |
| STEM121 | Y40410 | Takara Bio | 1:100 |
| Anti-Human Nuclei | MAB1281 | Sigma-Aldrich | 1:300 |
| Synaptophysin | ab8049 | Abcam | 1:50 |
| Rhodopsin | MA1-722 | Invitrogen | 1:100 |
| ZO-1 | 33-9100 | Invitrogen | 1:500 |
| alpha acetil tub | sc-23950 | Santa Cruz | 1:200 |
| PKC-a | SC-8393 | Santa Cruz | 1:200 |
| PCNA | MA5-11358 | Invitrogen | 1:100 |
| CD8 alpha | NB100-65729 | Novus | 1:200 |
| CD4 | 07-0403 | Invitrogen | 1:200 |
| CtBP2 (Ribbon) | PA5-79085 | Invitrogen | 1:300 |
| Ki67 | ab15580 | Abcam | 1:1000 |
| GFAP | Z0334 | Dako | 1:250 |
| SOX2 | ab97959 | Abcam | 1:1000 |
| MAP2 | ab5392 | Abcam | 1:250 |
| Phalloidin | | | |
| S-Opsin | 600-101-MP7 | Rochland | 1:200 |
| mGluR6 | NB300-189 | Novus | 1:100 |
| Synapsin | sc-376623 | Santa Cruz | 1:50-1:500 |
| PSD-95 | MA1-046 | Invitrogen | 1:20 |
| Syntaxin | ab41453 | Abcam | 1ug/ml |
