## Supplementary Figures for "Sub-retinal transplantation of human iPSC-derived retinal sheets: A promising approach for the treatment of macular degeneration"

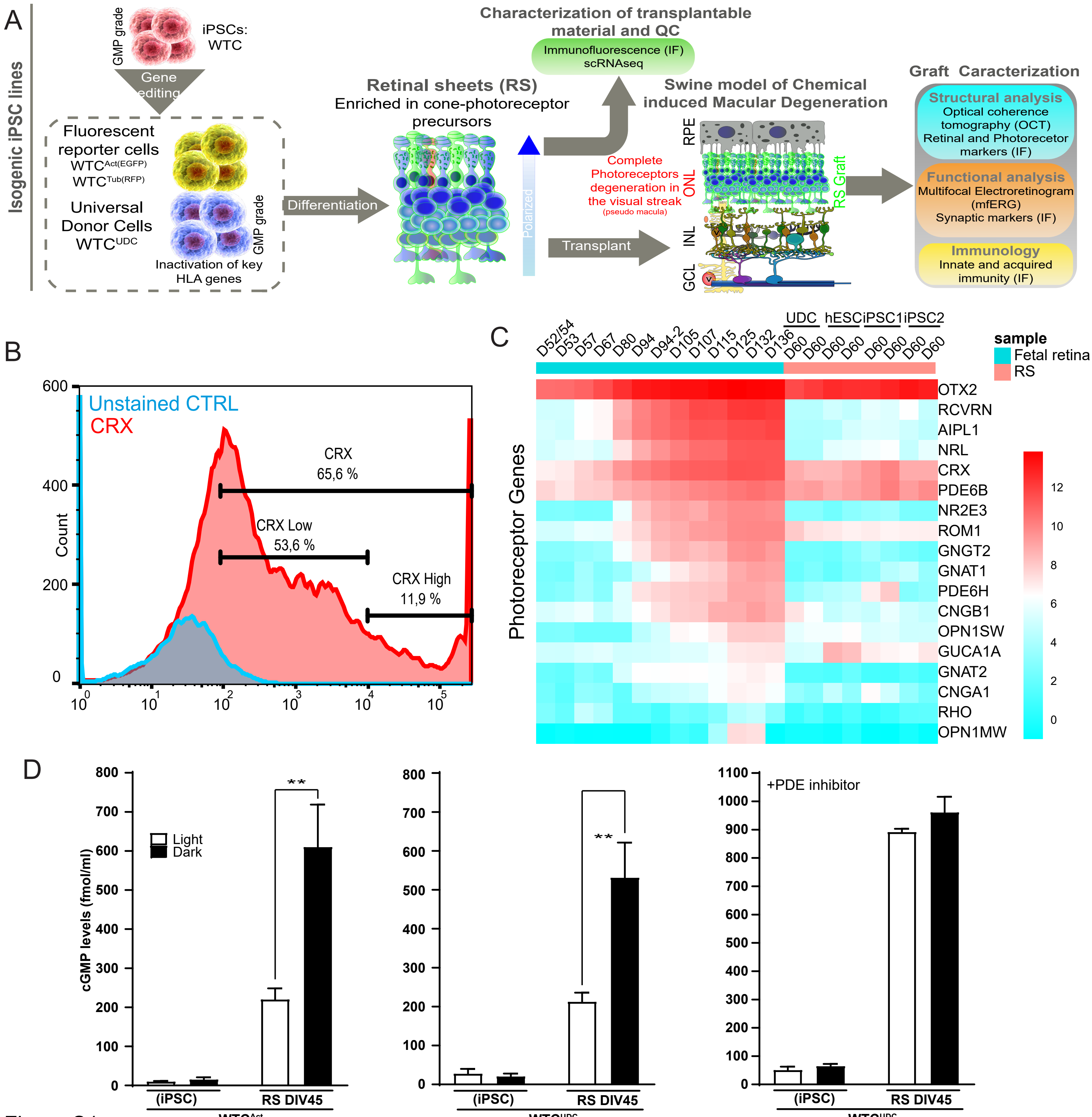
A
Isogenic iPSC lines
GMP grade
iPSCs: WTC
Gene editing
Fluorescent reporter cells
WTC Act(EGFP)
WTC Tub(RFP)
Universal Donor Cells
WTC UDC
GMP grade
Inactivation of key HLA genes
Differentiation
Characterization of transplantable material and QC
Immunofluorescence (IF)
scRNAseq
Retinal sheets (RS)
Enriched in cone-photoreceptor precursors
Polarized
Transplant
Swine model of Chemical induced Macular Degeneration
Complete Photoreceptors degeneration in the visual streak
(pseudo macula)
RPE
ONL
INL
GCL
RS Graft
Graft Caracterization
Structural analysis
Optical coherence tomography (OCT)
Retinal and Photorecetor markers (IF)
Functional analysis
Multifocal Electroretinogram (mfERG)
Synaptic markers (IF)
Immunology
Innate and acquired immunity (IF)
B
Unstained CTRL
CRX
CRX 65,6 %
CRX Low 53,6 %
CRX High 11,9 %
Count
600
400
200
0
10^0
10^1
10^2
10^3
10^4
10^5
C
Photoreceptor Genes
D52/54 D53 D57 D67 D80 D94 D94-2 D105 D107 D115 D125 D132 D136
UDC hESC iPSC1 iPSC2
D60 D60 D60 D60 D60 D60 D60 D60
sample
Fetal retina
RS
OTX2
RCVRN
AIPL1
NRL
CRX
PDE6B
NR2E3
ROM1
GNGT2
GNAT1
PDE6H
CNGB1
OPN1SW
GUCA1A
GNAT2
CNGA1
RHO
OPN1MW
12
10
8
6
4
2
0
D
cGMP levels (fmol/ml)
Light
Dark
**
+PDE inhibitor
(iPSC)
RS DIV45
WTC Act
WTC UDC


Figure S1

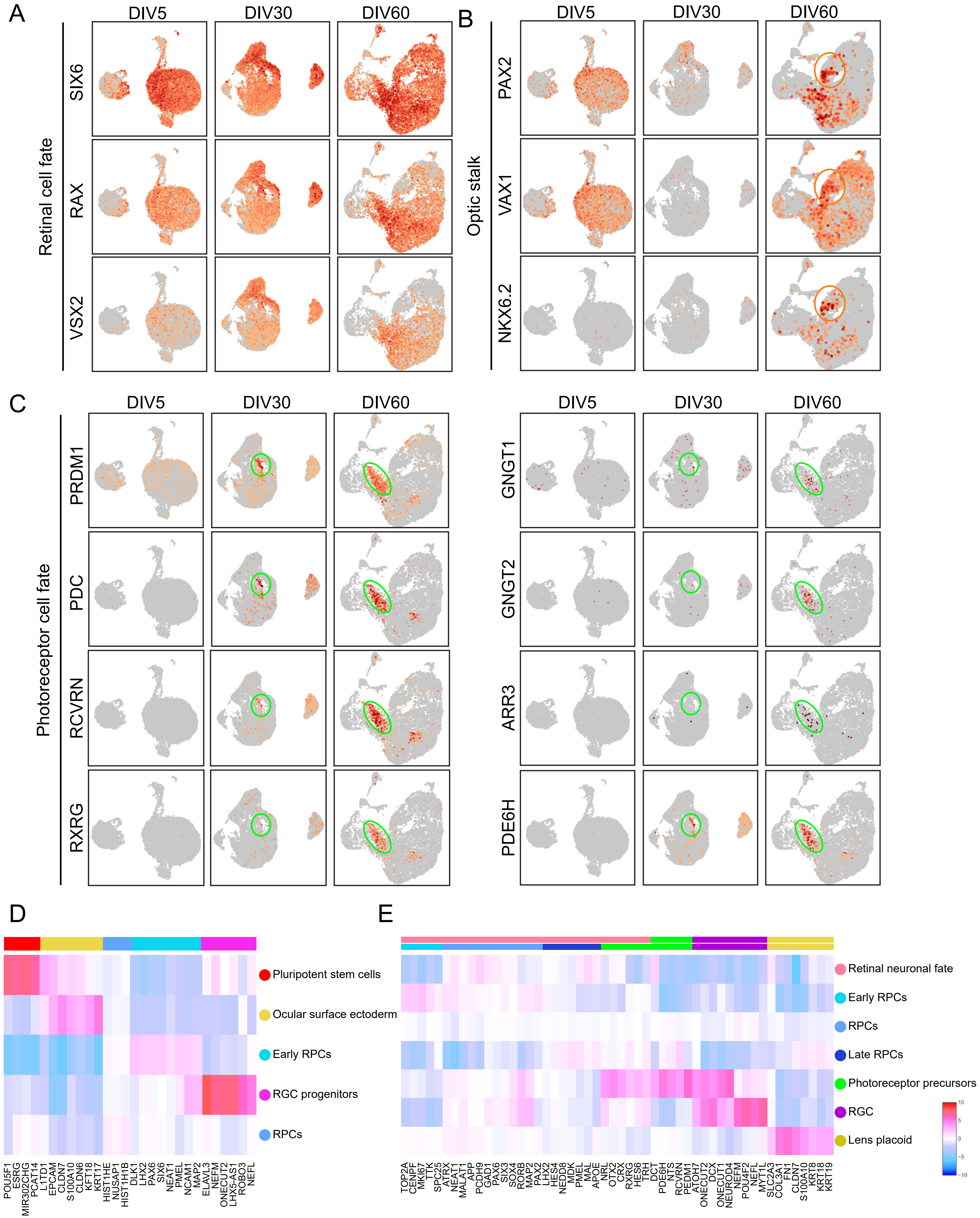

A
DIV5
DIV30
DIV60
Retinal cell fate
SIX6
RAX
VSX2
B
DIV5
DIV30
DIV60
Optic stalk
PAX2
VAX1
NKX6.2
C
DIV5
DIV30
DIV60
Photoreceptor cell fate
PRDM1
PDC
RCVRN
RXRG
DIV5
DIV30
DIV60
GNGT1
GNGT2
ARR3
PDE6H
D
Pluripotent stem cells
Ocular surface ectoderm
Early RPCs
RGC progenitors
RPCs
POU5F1
ESRG
MIR302CHG
PCAT14
L1TD1
EPCAM
CLDN7
S100A10
CLDN6
KFT18
KRT17
HIST1HE
NUSAP1
HIST1H1B
DLK1
LHX2
PAX6
SIX6
NEAT1
PMEL
NCAM1
MAP2
ELAVL3
NEFM
ONECUT2
LHX5-AS1
ROBO3
NEFL
E
Retinal neuronal fate
Early RPCs
RPCs
Late RPCs
Photoreceptor precursors
RGC
Lens placoid
10
5
0
-5
-10
TOP2A
CENPF
MKI67
TTK
SPC25
ATRX
NEAT1
MALAT1
APP
PCDH9
GAD1
PAX6
SIX3
SOX4
RORB
MAP2
PAX2
LHX2
HES4
NEDD8
MDK
PMEL
MAL
APOE
NRL
OTX2
CRX
RXRG
HES6
TRH
DCT
PDE6H
NTS
RCVRN
PEDM1
ATOH7
ONECUT2
DCX
ONECUT1
NEUROD4
NEFM
POU4F2
NEFL
MYT1L
SLC2A3
COL3A1
FN1
CLDN7
S100A10
KRT8
KRT18
KRT19


Figure S2

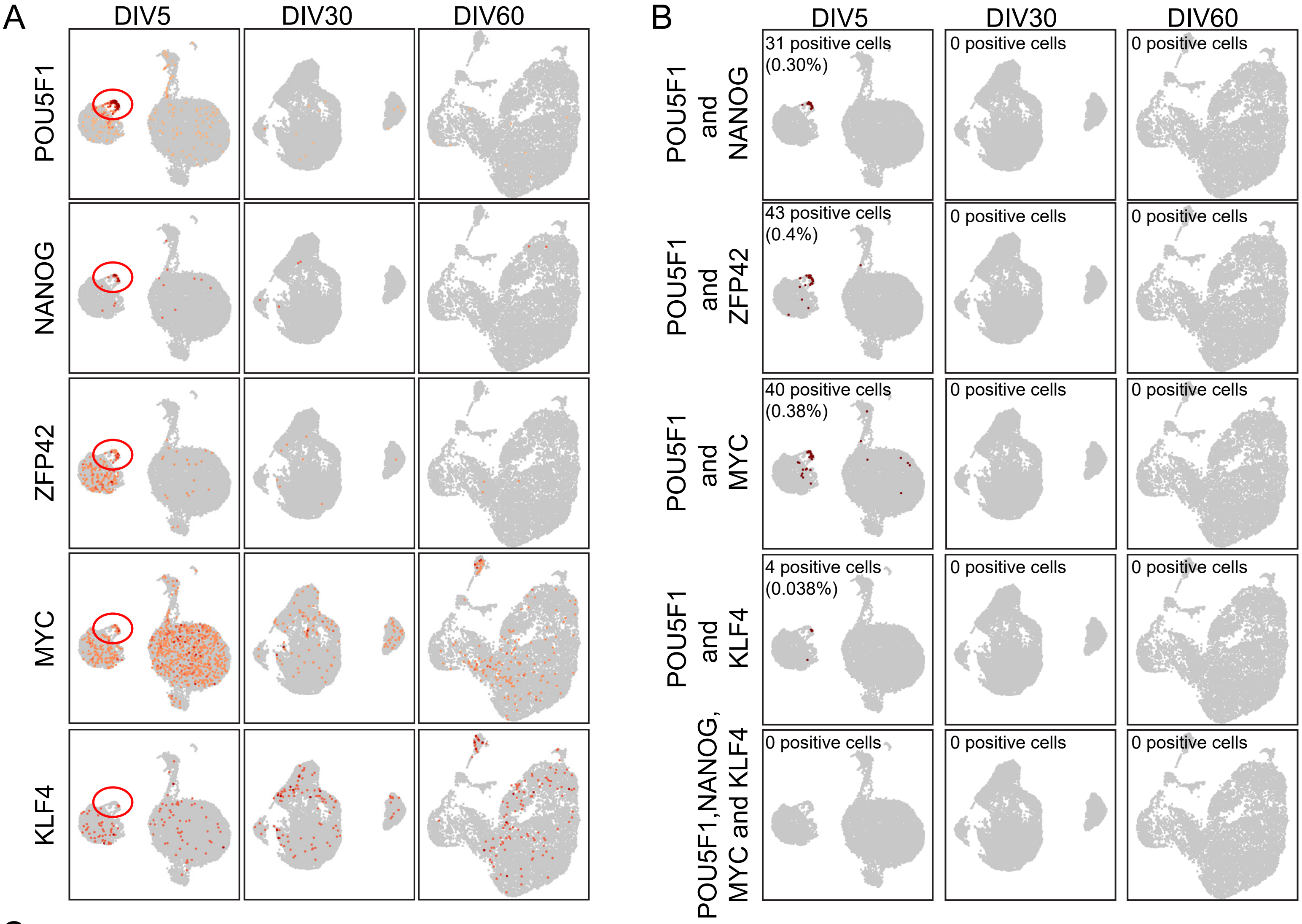


C

**Summary**

| cell type | number of cells injected | number injection | number of developping teratoma | observation of last teratoma (day after injection) | % of teratoma formation |
|---|---|---|---|---|---|
| ESCs | $2.5x10^6$ | 4 | 3 | 44 | 75% |
| iPSCs | $2.5x10^6$ | 8 | 8 | 26 | 100% |
| ESC-derived RSs | $2.5x10^6$ | 4 | 0 | \ | 0% |
| iPSC-derived RSc | $2.5x10^6$ | 22 | 0 | \ | 0% |

Figure S3

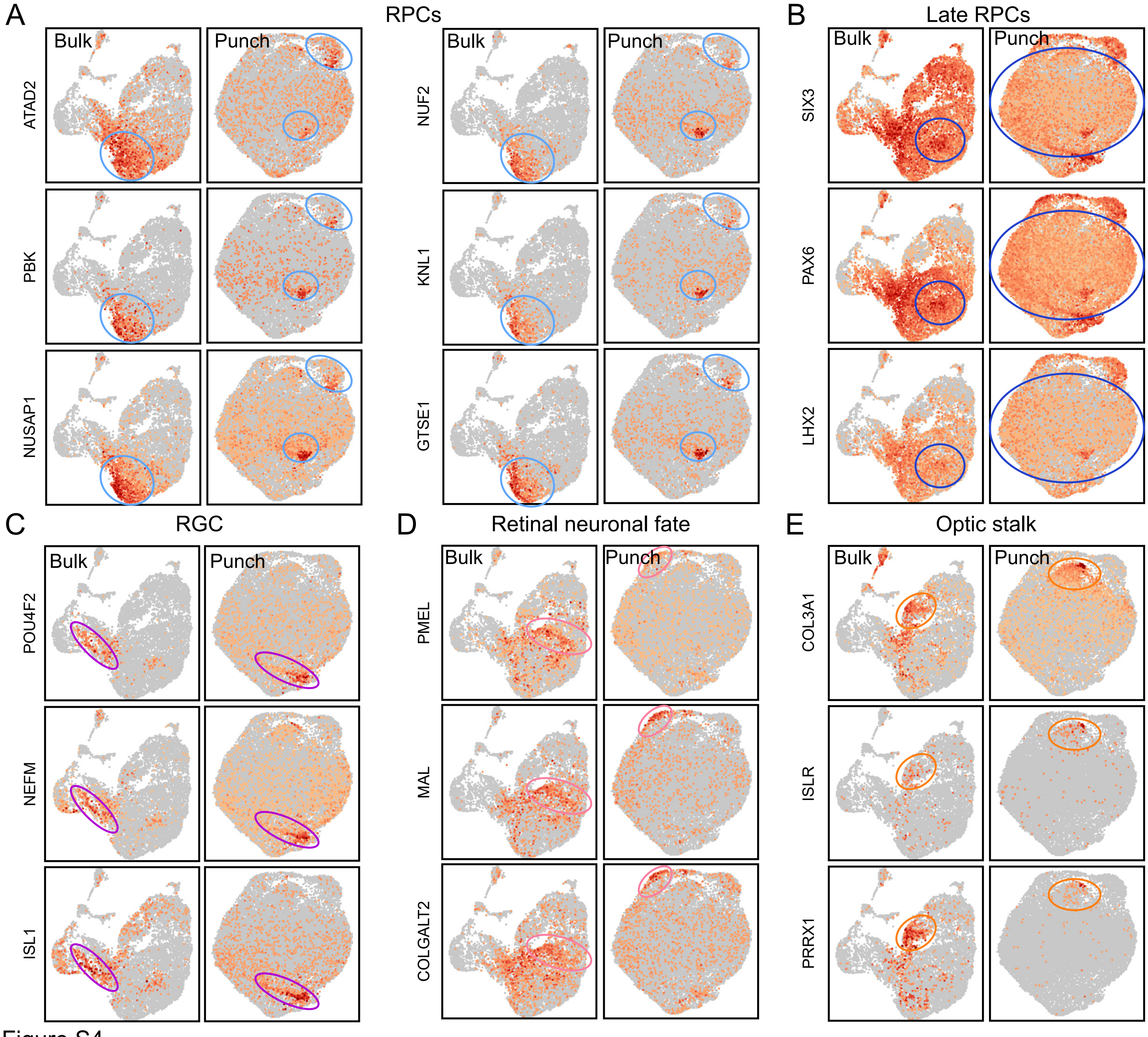
A
RPCs
Bulk
Punch
ATAD2
PBK
NUSAP1
NUF2
KNL1
GTSE1
B
Late RPCs
SIX3
PAX6
LHX2
C
RGC
POU4F2
NEFM
ISL1
D
Retinal neuronal fate
PMEL
MAL
COLGALT2
E
Optic stalk
COL3A1
ISLR
PRRX1

Figure S4

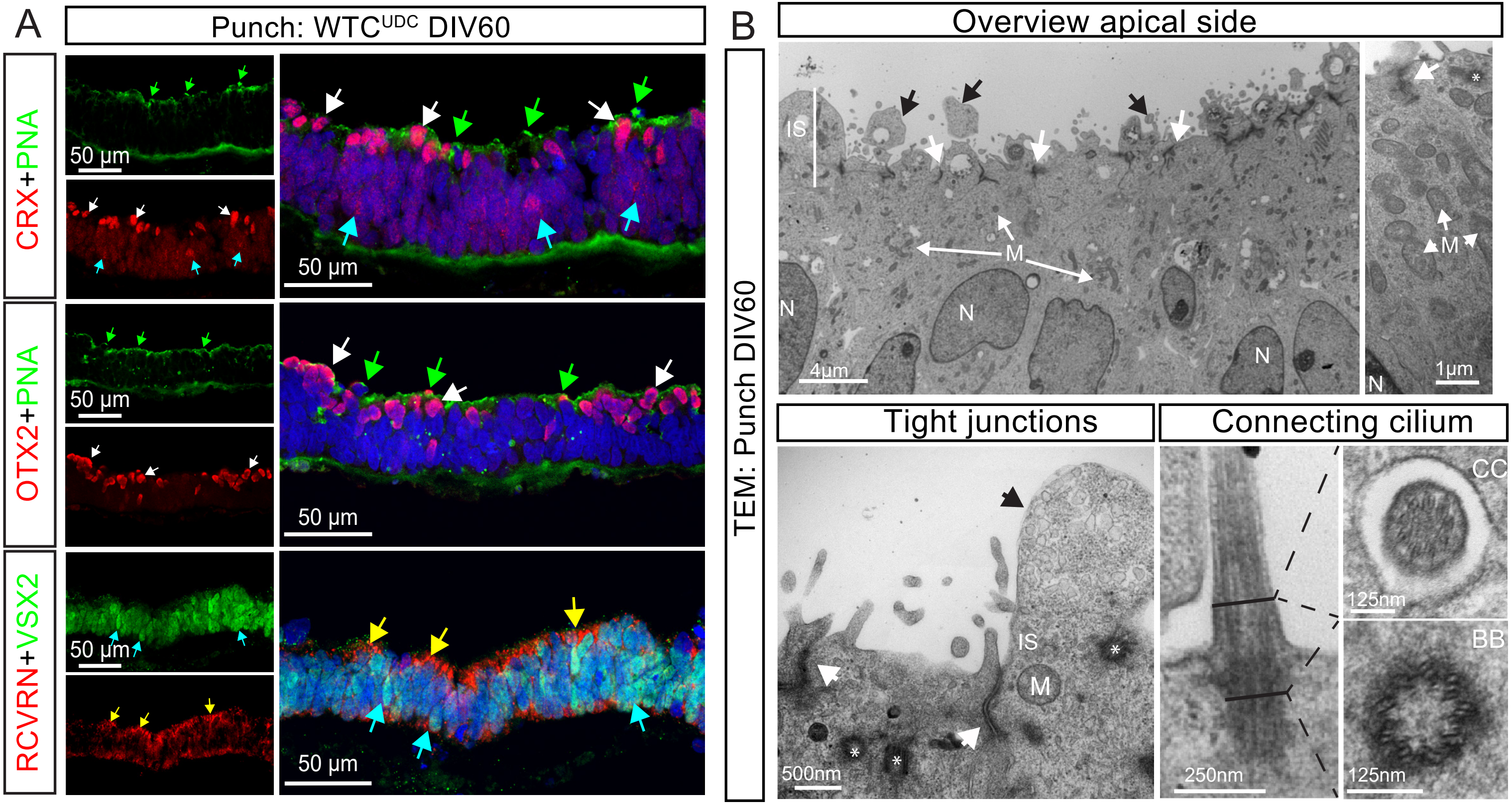
A
Punch: WTC UDC DIV60
CRX+PNA
OTX2+PNA
RCVRN+VSX2
50 µm
B
TEM: Punch DIV60
Overview apical side
IS
M
N
4µm
1µm
Tight junctions
Connecting cilium
500nm
250nm
125nm
CC
BB

Figure S5

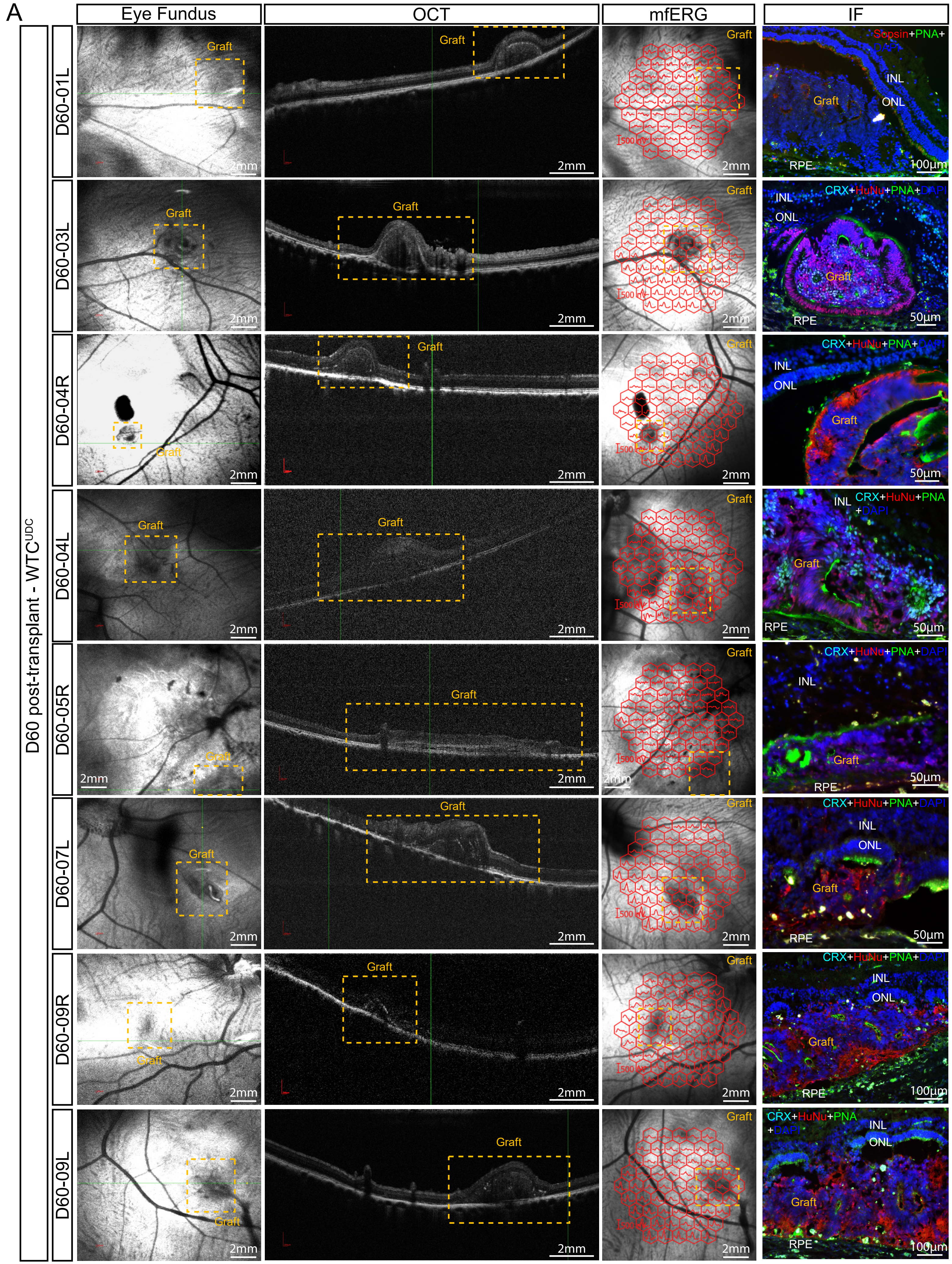
A
Eye Fundus
OCT
mfERG
IF
D60 post-transplant - WTC UDC
D60-01L
D60-03L
D60-04R
D60-04L
D60-05R
D60-07L
D60-09R
D60-09L
Graft
2mm
500 nV
INL
ONL
RPE
100µm
50µm
Sopsin+PNA+DAPI
CRX+HuNu+PNA+DAPI

Figure S6

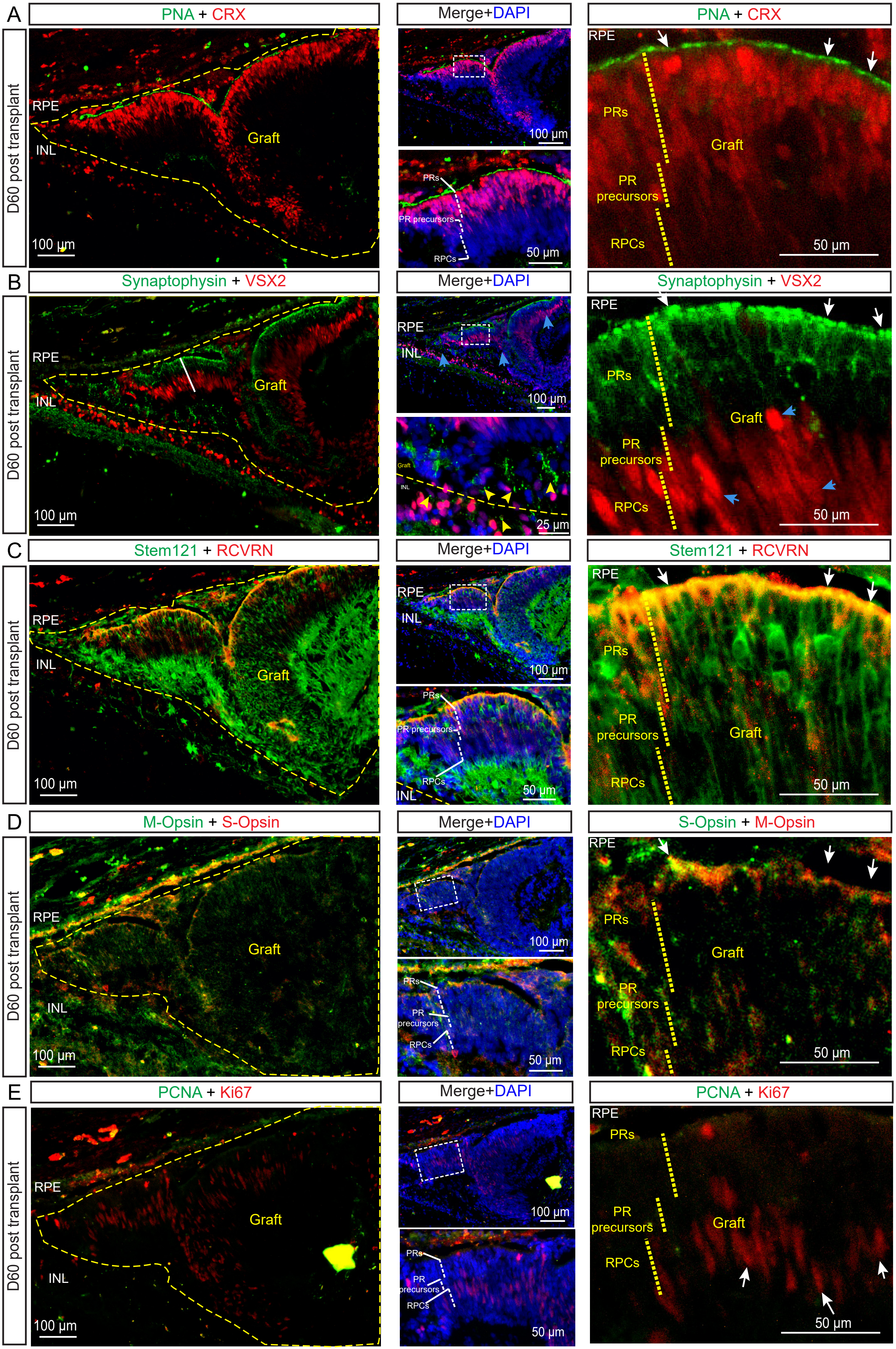

A
PNA + CRX
Merge+DAPI
PNA + CRX
D60 post transplant
RPE
INL
Graft
100 µm
100 µm
PRs
PR precursors
RPCs
50 µm
RPE
PRs
Graft
PR
precursors
RPCs
50 µm
B
Synaptophysin + VSX2
Merge+DAPI
Synaptophysin + VSX2
D60 post transplant
RPE
INL
Graft
100 µm
RPE
INL
100 µm
Graft
INL
25 µm
RPE
PRs
Graft
PR
precursors
RPCs
50 µm
C
Stem121 + RCVRN
Merge+DAPI
Stem121 + RCVRN
D60 post transplant
RPE
INL
Graft
100 µm
RPE
INL
100 µm
PRs
PR precursors
RPCs
INL
50 µm
RPE
PRs
PR
precursors
Graft
RPCs
50 µm
D
M-Opsin + S-Opsin
Merge+DAPI
S-Opsin + M-Opsin
D60 post transplant
RPE
Graft
INL
100 µm
100 µm
PRs
PR
precursors
RPCs
50 µm
RPE
PRs
Graft
PR
precursors
RPCs
50 µm
E
PCNA + Ki67
Merge+DAPI
PCNA + Ki67
D60 post transplant
RPE
Graft
INL
100 µm
100 µm
PRs
PR
precursors
RPCs
50 µm
RPE
PRs
PR
precursors
Graft
RPCs
50 µm


Figure S7

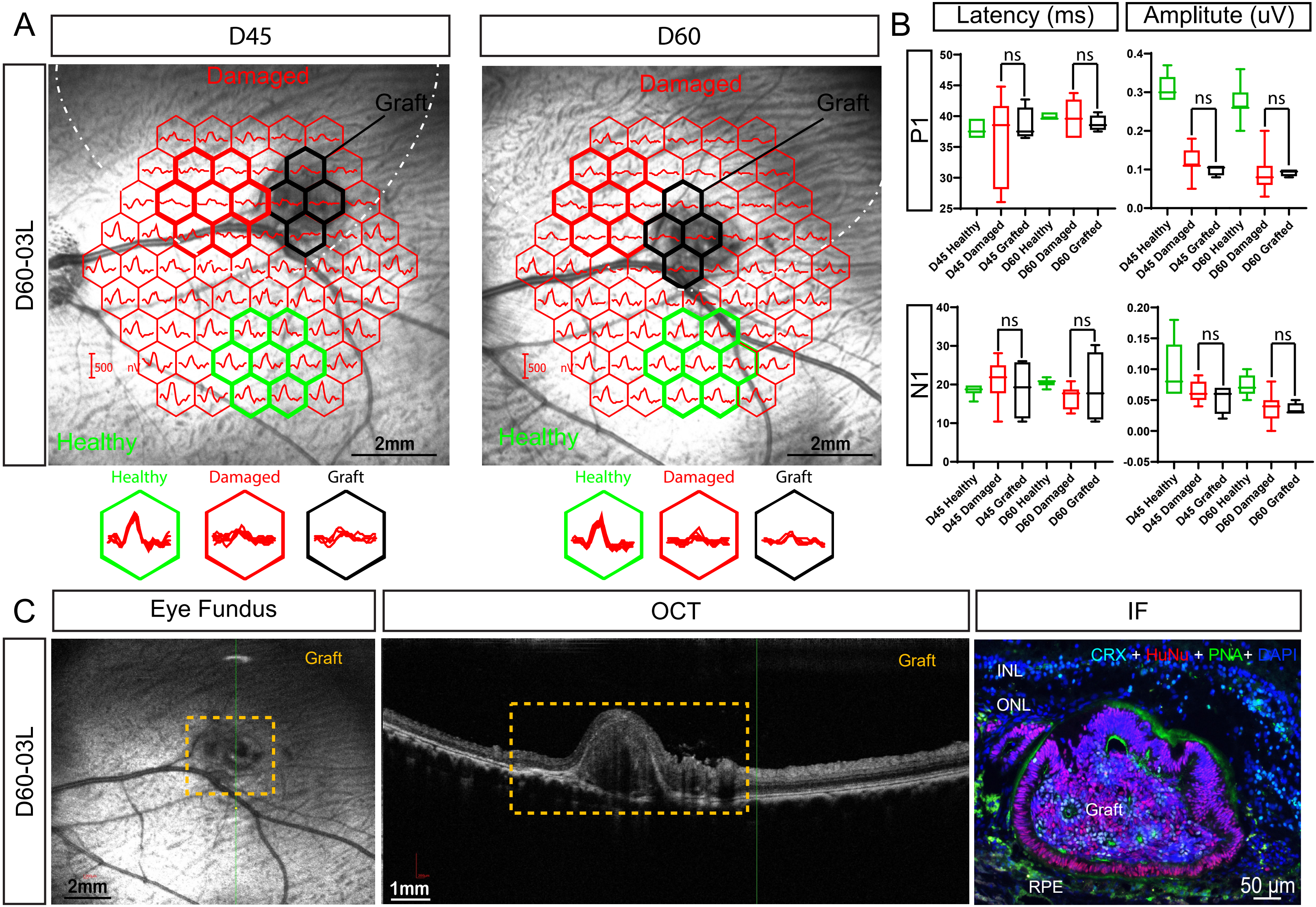

A
D45
D60
D60-03L
Damaged
Graft
Healthy
500 nV
2mm
Healthy
Damaged
Graft
B
Latency (ms)
Amplitute (uV)
P1
N1
ns
D45 Healthy
D45 Damaged
D45 Grafted
D60 Healthy
D60 Damaged
D60 Grafted
C
Eye Fundus
OCT
IF
Graft
1mm
CRX + HuNu + PNA+ DAPI
INL
ONL
RPE
50 µm


Figure S8

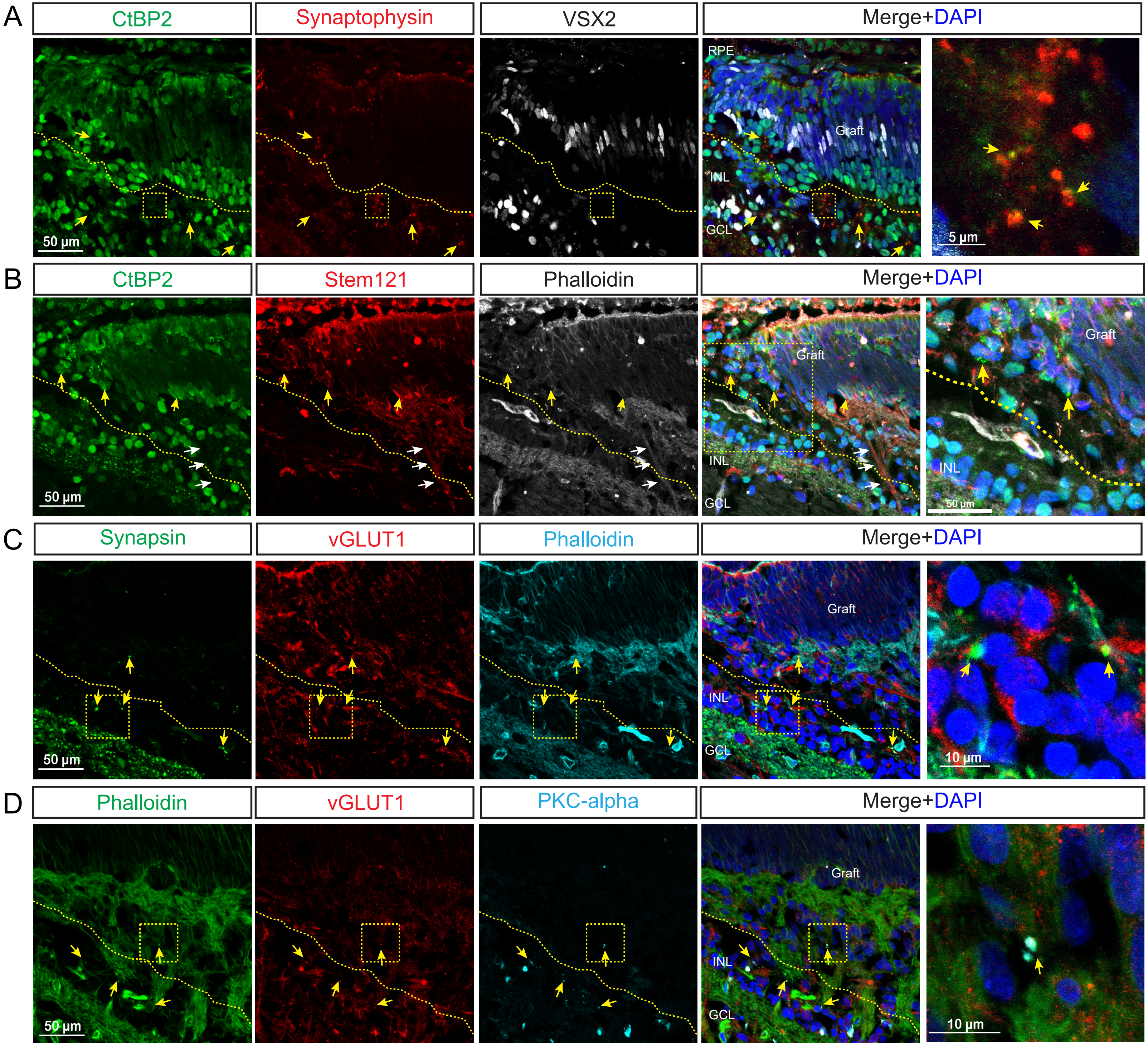
A
CtBP2
Synaptophysin
VSX2
Merge+DAPI
RPE
Graft
INL
GCL
50 µm
5 µm
B
CtBP2
Stem121
Phalloidin
Merge+DAPI
Graft
INL
GCL
50 µm
C
Synapsin
vGLUT1
Phalloidin
Merge+DAPI
Graft
INL
GCL
50 µm
10 µm
D
Phalloidin
vGLUT1
PKC-alpha
Merge+DAPI
Graft
INL
GCL
50 µm
10 µm

Figure S9

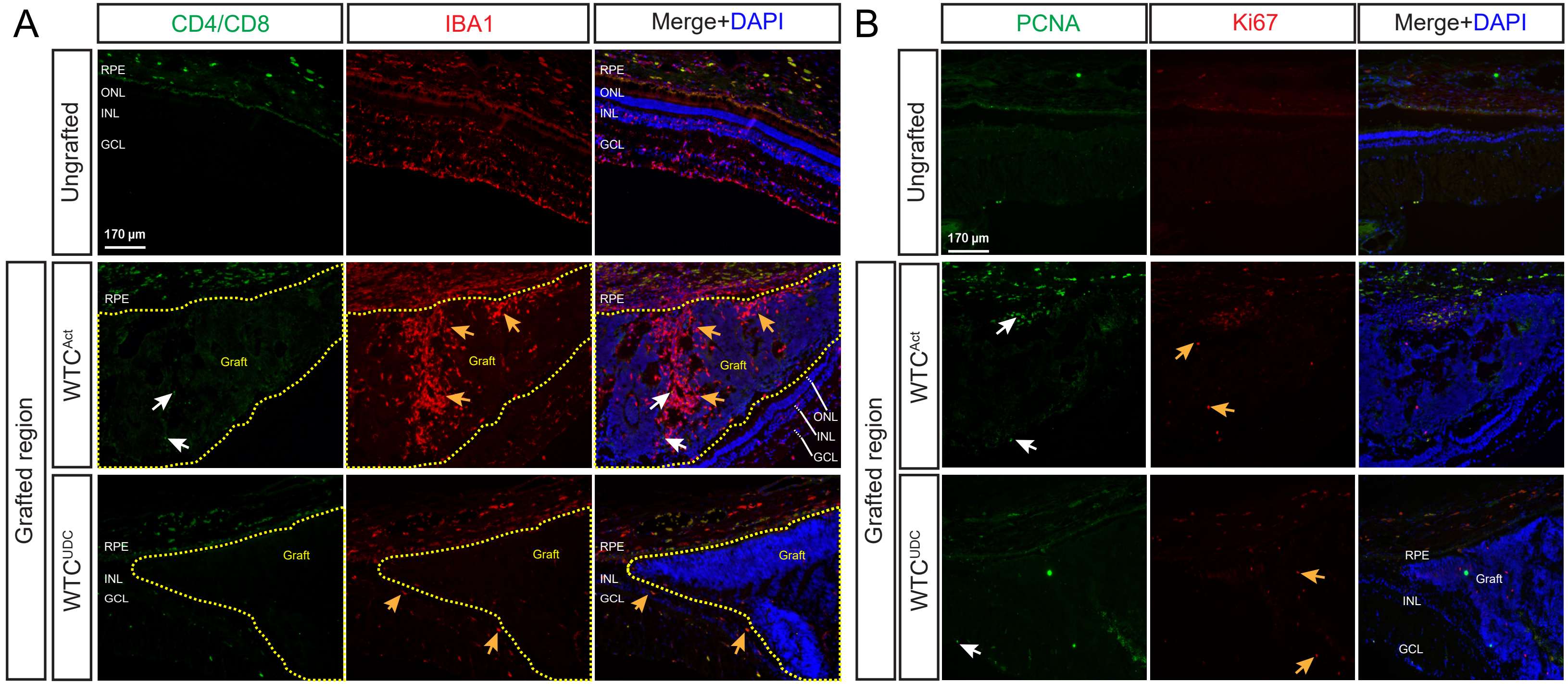


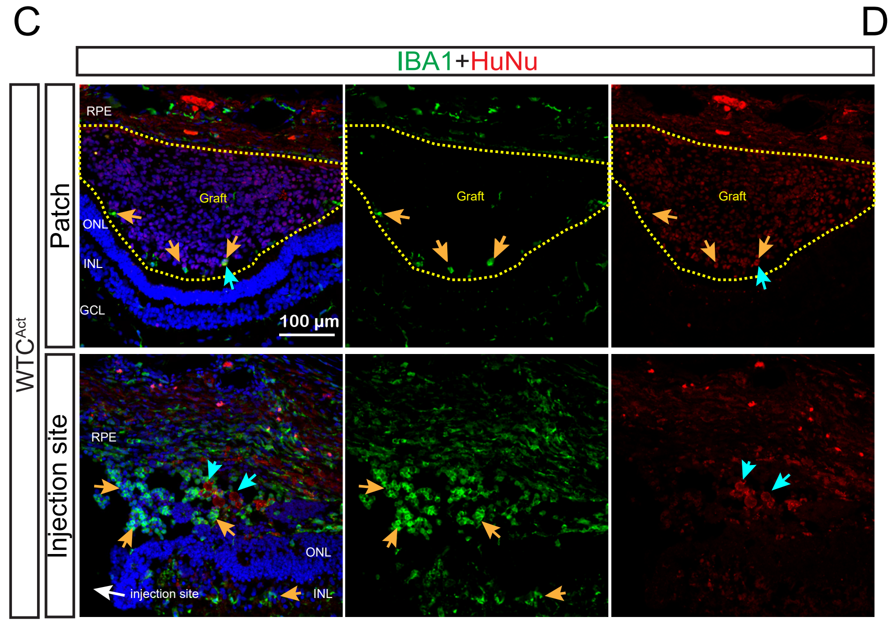


D

| Animal | Cell line | Eye | Immune response in the lesion | | Immune response on graft | | Proliferation |
|---|---|---|---|---|---|---|---|
| D60-01 | WTC^UDC | Right | IBA1 | CD4/CD8 | IBA1 | CD4/CD8 | Ki67/PCNA |
| | | Left | Low | Few Mϕ IBA1+/CD4+ | Low | NO | Low |
| D60-03 | | Right | Low | Few Mϕ IBA1+/CD4+ | Low | NO | Low |
| | | Left | Low | NO | Low | NO | Low |
| D60-04 | | Right | High | Few Mϕ IBA1+/CD4+ | High | NO | Low |
| | | Left | Low | NO | Low | NO | Low |
| D60-05 | | Right | Low | NO | Low | NO | Low |
| | | Left | NA | NA | NA | NA | NA |
| D60-07 | | Right | NA | NA | NA | NA | NA |
| | | Left | Low | Few Mϕ IBA1+/CD4+ | Low | NO | Low |
| D60-09 | | Right | Low | Few Mϕ IBA1+/CD4+ | Low | NO | Low |
| | | Left | Low | Few Mϕ IBA1+/CD4+ | Low | NO | Low |
| D60-12 | WTC^Act | Right | NA | NA | NA | NA | NA |
| | | Left | High | NO | High | NO | Low |
| D60-13 | | Right | High | NO | Very High | NO | Low |
| | | Left | NA | NA | NA | NA | NA |
| D60-15 | | Right | High | NO | High | NO | Low |
| | | Left | High | NO | High | NO | Low |
| D60-17 | | Right | NA | NA | NA | NA | NA |
| | | Left | High | NO | High | NO | Low |

Figure S10

| Cell line | Animal | Eye | Lesion | Transplant | Number of punchs | Cells | Punch size |
|---|---|---|---|---|---|---|---|
| $WTC^{UDC}$ | D30-07 | Right | PERFECT | Success (no visual) | 1 | $WTC^{UDC}$ | 3 mm |
| | | Left | PERFECT | Failed | NA | NA | NA |
| | D60-01 | Right | PERFECT | Failed | NA | NA | NA |
| | | Left | PERFECT | Success | 1 | $WTC^{UDC}$ | 3 mm |
| | D60-02 | Right | NO | NA | NA | NA | NA |
| | | Left | PERFECT | Failed | NA | NA | NA |
| | D60-03 | Right | GOOD (SEVERE) | Success | 1 | $WTC^{UDC}$ | 3 mm |
| | | Left | GOOD (SEVERE) | Success | 1 | $WTC^{UDC}$ | 3 mm |
| | D60-04 | Right | PERFECT | Success | 1 | $WTC^{UDC}$ | 2 mm |
| | | Left | GOOD (SEVERE) | Success | 2 | $WTC^{UDC}$ | 2 + 3 mm |
| | D60-05 | Right | PERFECT | Success | 2 | $WTC^{UDC}$ | 2 mm |
| | | Left | GOOD (SEVERE) | Success | 2 | $WTC^{UDC}$ | 2 mm |
| | D60-06 | Right | GOOD (SUBTILE) | Success | 2 | $WTC^{UDC}$ | 2 mm |
| | | Left | NO | Success | 6 | $WTC^{UDC}$ | 1 mm |
| | D60-07 | Right | NO | NA | NA | NA | NA |
| | | Left | GOOD (SEVERE) | Success | 2 | $WTC^{UDC}$ | 2 mm |
| | D60-08 | Right | NO | NA | NA | NA | NA |
| | | Left | GOOD (SUBTILE) | Success | 4 | $WTC^{UDC}$ | 1.5 mm |
| | D60-09 | Right | PERFECT | Success | 3 | $WTC^{UDC}$ | 2 mm |
| | | Left | PERFECT | Success | 1 | $WTC^{UDC}$ | 2 mm |
| | D60-10 | Right | PERFECT | Success | 3 | $WTC^{UDC}$ | 1.5 mm |
| | | Left | NO | NA | NA | NA | NA |
| | D60-11 | Right | PERFECT | Success | 7 | $WTC^{UDC}$ | 1.5 mm |
| | | Left | NO | NA | NA | NA | NA |
| $WTC^{Act}$ | D60-12 | Right | NO | NA | NA | NA | NA |
| | | Left | PERFECT | Success | 3 | $WTC^{Act}$ | 2 mm |
| | D60-13 | Right | GOOD (SUBTILE) | Success | 5 | $WTC^{Act}$ | 2 mm |
| | | Left | NO | NA | NA | NA | NA |
| | D60-14 | Right | Died after cobalt chloride injection | | | | |
| | | Left | | | | | |
| | D60-15 | Right | GOOD (SEVERE) | Success | 2 | $WTC^{Act}$ | 3 mm |
| | | Left | GOOD (SEVERE) | Success | 2 | $WTC^{Act}$ | 2 mm |
| | D60-16 | Right | PERFECT | Success | 2 | $WTC^{Act}$ | 2 mm |
| | | Left | GOOD (SEVERE) | Failed | NA | NA | NA |
| | D60-17 | Right | PERFECT | Success | 2 | $WTC^{Act}$ | 2 mm |
| | | Left | PERFECT | Success | 2 | $WTC^{Act}$ | 2 mm |

Table S1

| Cell line | Animal | Eye | OCT/ERG D30 | OCT/ERG D45 | OCT/ERG D60 | Graft: Inside the lesion | Graft: Flat areas | Graft: Polarisation | CRX | PNA | SCORE |
|---|---|---|---|---|---|---|---|---|---|---|---|
| $WTC^{UDC}$ | D60-01 | Right | NA | NA | NA | NA | NA | NA | NA | NA | 0/10 |
| | | Left | NA | NA | NA | NO | NO | Partially | YES | YES | 6/10 |
| | D60-03 | Right | Eye mouvement, no analysis | YES | YES | YES | YES | YES | YES | YES | 10/10 |
| | | Left | YES | NO | NO | Partially | NO | Partially | Rosettes only | Rosettes only | 5/10 |
| | D60-04 | Right | YES | YES | YES | NO | YES | YES | Rosettes only | Rosettes only | 7/10 |
| | | Left | Eye mouvement, no analysis | YES | Cataract | Partially | Partially | Partially | Rosettes only | Rosettes only | 7/10 |
| | D60-05 | Right | NO | NO | NA | YES | Partially | NO | LOW | LOW | 4/10 |
| | | Left | Kenalog shadow, no analysis | Cataract | NA | NA | NA | NA | NA | NA | 0/10 |
| | D60-07 | Right | NA | NA | NA | NA | NA | NA | NA | NA | 0/10 |
| | | Left | NO | NA | YES | NO | NO | NO | LOW | LOW | 2/10 |
| | D60-09 | Right | NO | NA | YES | Partially | Partially | Partially | Rosettes only | Rosettes only | 6/10 |
| | | Left | NA | NA | NO | Partially | NO | Partially | Rosettes only | Rosettes only | 5/10 |
| $WTC^{Act}$ | D60-12 | Right | NA | NA | NA | NA | NA | NA | NA | NA | 0/10 |
| | | Left | Kenalog shadow, no analysis | NA | Cataract | NO | NO | Partially | Rosettes only | NO | 2/10 |
| | D60-13 | Right | YES | NA | Cataract | NO | NO | NO | Graft damaged by immune cells | | 2/10 |
| | | Left | NA | NA | NA | NA | NA | NA | NA | NA | 0/10 |
| | D60-15 | Right | Eye mouvement, no analysis | NA | Cataract | NO | YES | NO | Graft damaged by immune cells | | 2/10 |
| | | Left | NO | NA | NO | NO | YES | NO | Graft damaged by immune cells | | 2/10 |
| | D60-16 | Right | NA | NA | NA | Partially | YES | NO | Graft damaged by immune cells | | 3/10 |
| | | Left | NA | NA | NA | NA | NA | NA | NA | NA | 0/10 |
| | D60-17 | Right | NA | NA | NA | Partially | YES | NO | Graft destroyed by immune cells | | 4/10 |
| | | Left | Cataract, no analysis | NA | Cataract | Partially | YES | Partially | Rosettes only | NO | 4/10 |

Table S2
